# Prp16 enables efficient splicing of introns with diverse exonic consensus elements in the short-intron rich *Cryptococcus neoformans* transcriptome

**DOI:** 10.1101/2024.12.05.626984

**Authors:** Manendra Singh Negi, Vishnu Priya Krishnan, Niharika Saraf, Usha Vijayraghavan

**Author notes:** Corresponding author’s.

## Abstract

The DEAH box splicing helicase Prp16 in budding yeast governs spliceosomal remodeling from the branching conformation (C complex) to the exon ligation conformation (C* complex). In this study, we examined the genome-wide functions of Prp16 in the short intron-rich genome of the basidiomycete yeast *Cryptococcus neoformans*. The presence of multiple introns per transcript with intronic features more similar to higher eukaryotes makes it a promising model to study spliceosomal splicing. Using a promoter-shutdown conditional Prp16 knockdown strain, we uncovered its genome-wide but substrate-specific roles in *C. neoformans* splicing. The splicing functions of Prp16 are dependent on its helicase motif I and motif II that are conserved motifs for helicase activity. A small subset of introns spliced independent of Prp16 activity, were investigated to discover that exonic sequences at the 5’ splice site (5’SS) and 3’ splice site (3’SS) with stronger affinity for U5 loop 1 as a common feature in these introns. Furthermore, short (60-100nts) and ultra-short introns (<60nts) prevalent in the *C. neoformans* transcriptome were more sensitive to Prp16 knockdown than longer introns, indicating Prp16 is required for the efficient splicing of short and ultra-short introns. We propose that stronger U5 snRNA-pre-mRNA interactions enable the efficient transition of the spliceosome from the first to the second catalytic confirmation in Prp16 knockdown, particularly for short introns and introns with suboptimal features. This study provides insights into the fine-tuning spliceosomal helicase functions with variations in *cis-*element features.

## Introduction

Removal of introns and splicing of exons of RNA pol II transcribed pre-mRNA is ubiquitous in eukaryotic gene expression. This process is executed by a versatile multimegadalton complex composed of 5 snRNPs (U1, U2, U4, U5, and U6) and over 150 proteins known as spliceosomes [1]. *In vitro* splicing assay has been key to deciphering the stepwise assembly of spliceosomal components on model yeast/ human pre-mRNAs. The systematic assembly by conformational and compositional remodeling ensures the two sequential transesterification reactions: first, the branching leading to lariat intermediate and then exon ligation [2]. The characteristic conserved *cis* sequences at the 5’ splice site (5’SS), branch point sequence (BPS) and 3’ splice site (3’SS) and their interaction with spliceosomal snRNAs guide spliceosome assembly, activation and catalysis [3]. The compositional and conformational remodeling is facilitated by 8 ATP-dependent RNA helicases that disrupt or translocate various RNA-protein, RNA-RNA and protein-protein interactions [4]. During *in vitro* spliceosome assembly, budding yeast DEAD-Box helicases Sub2/UAP56, Prp5, and Prp28 ensure accurate splice site recognition to form the B complex. The Brr2 helicase unwinds the U4/U6 duplex to release U4, during activation of the B complex, which is followed by the recruitment of NTC to form the B^act^ complex. DEAH box helicase Prp2 remodels the B^act^ complex to the catalytically active B* complex wherein the first catalytic reaction generates the intermediates 5’exon and lariat intron-3’exon held in the C complex. The Prp16 DEAH box helicase remodels the C complex by destabilizing the binding of branching factors like Cwc25 and allows the recruitment of exon ligation factors such as Slu7, Prp18 thus, forming C* complex primed for exon ligation. The second catalytic reaction ligates 5’ and 3’ exons and excises the lariat intron. The release of the spliced mRNA from the post-spliceosomal (P) complex is mediated by Prp22-helicase, while the Prp43-helicase mediates intron lariat spliceosome (ILS) disassembly [5].

A suppressor mutant screen of *cis* intronic invariant branch nucleotide A to C mutation in a budding yeast splicing reporter minigene identified Prp16 [6,7] as factor important for branch recognition. Subsequent other work with mutants with diminished ATPase activity also showed aberrant use of pre-mRNAs with mutant branch nucleotide C. Thus, it was inferred that the ATPase activity of Prp16 ensures splicing fidelity [8]. Subsequent *in vitro* splicing assays showed an important role for *S. cerevisiae* Prp16-mediated proofreading of substrate before first step catalysis [9]. Prp16 mutants defective for ATPase activity stabilized the interaction of first step splicing factors Yju2 and Cwc25 with the mutated substrate, thereby supporting progression to form the branched lariat intron-3’exon. However, the ATPase activity of Prp16 destabilizes the branching factors by remodeling spliceosome after 1^st^ step catalysis to facilitate exon ligation [10,11]. This dual function of Prp16 leads to a hypothesis where the ATPase activity of Prp16 competes with its ATP-independent facilitation of 5’SS cleavage at the first step to proofread slow branching reaction [12].

The *in vivo* genome wide relevance of Prp16-mediated spliceosome remodeling on the global transcriptome has not been explored much. In fission yeast, Prp16 is required for the splicing of a vast majority of the introns in its genome where its action facilitates first step catalysis by destabilizing 5’SS-U6snRNA and BS-U2snRNA interactions [13]. Further, *Sp*Prp16 functions impact fission yeast cell cycle progression and heterochromatinization of centromeres and telomeres [13,14]. In many species with transcriptomes enriched with introns, the diversity of the *cis*-elements and their interactions with spliceosomal factors machinery can play a critical role in splicing efficiency. Thus, these interactions and the kinetics of splicing progression add an additional layer of complexity to the dynamic regulation of gene expression [15]. This raises intriguing questions on how splicing factors, including snRNPs, accommodate the diversity of intronic *cis*-elements in a genome and across diverse genomes. While splicing factors and spliceosome assembly have been extensively studied in intron-poor yeast models such as *Saccharomyces cerevisiae* and, to a lesser extent, in *Schizosaccharomyces pombe*, their role in intron-rich transcripts of *Cryptococcus neoformans* where over 99% of transcripts bear multiple introns remains relatively understudied. Notably, the exon-intron architecture of *C. neoformans,* with degenerate splice signals and variable intronic features, differs from the well-studied fungal models and resemble to intronic features found in higher eukaryotes (e.g., plants and humans). Therefore *C. neoformans* is a promising model for investigating splicing regulation in complex transcriptomes [16].

Here, we studied genome-wide splicing functions of *C. neoformans* Prp16 by generating a promoter-shutdown conditional knockdown strain. We uncovered a genome-wide but intron substrate-specific role of Prp16 for splicing *C. neoformans* introns. The intron-specific functions are in part contributed by pre-mRNA and U5 loop 1 interactions, which could be related to its role in remodeling the spliceosome for catalysis. Additionally, we report that the splicing of short and ultra-short introns is more sensitive to Prp16 depletion compared to longer introns, suggesting that Prp16 plays a crucial role in processing these specific types of introns.

## Results

### *C. neoformans* Prp16 is essential for growth and viability

*S. cerevisiae* Prp16 is an essential spliceosomal DEAH box helicase that carries out remodeling from the spliceosomal branching conformation formed after first-step catalysis (C complex) to second-step exon ligation conformation (C* complex) as deciphered from extensive *in vitro* splicing assays using model actin mini-transcripts. We identified the Prp16 ortholog in *C. neoformans* as CNAG_02303 using homology-based iterative HMMER [17] jackhammer search with *S. pombe* Prp16 as the input sequence. The identified *C. neoformans* Prp16 ortholog shares significant conservation at the C-terminal helicase domain with all six signatory motifs seen in other DEAD/H box helicases (Supplementary Figure 1a). These motifs are involved in ATP binding, hydrolysis and RNA duplex unwinding [13,18]. To explore the splicing functions for *C. neoformans* Prp16, we engineered a *C. neoformans* Prp16 conditional knockdown strain by swapping the native promoter with *Gal7* promoter and N-terminal mCherry tag (Figure 1(a)). The resulting strain, *PGAL7*:Prp16 would overexpress mCherry-Prp16 transcript when grown in permissive galactose media (YPG), and promoter shutdown would occur when grown in nonpermissive glucose (YPD) media (Figure 1(a), Supplementary Figure S2(a)). We tested Prp16 transcript levels and the mCherry-Prp16 protein level when grown in YPD media by RT-PCR, confocal microscopy and western blot (Supplementary Figure S2(b), S1(c), S1(d)). We standardized growth condition to achieve significant knockdown of Prp16 by growing primary culture in permissive YPG media followed by growth in nonpermissive YPD media for 12 hours and then followed by re-inoculation again this time into two cultures, one in nonpermissive media YPD and other in permissive media YPG conditions for 9 hours (Supplementary Figure S3(a)). In comparison with similarly treated wild-type (WT) cells, the growth kinetics of *PGAL7*:Prp16 strain was noticeably reduced after 9 hours of growth in nonpermissive YPD media. No significant growth alteration was observed between *PGAL7*:Prp16 and WT strains when grown in permissive YPG media (Figure 1(b), 1(c)). Similar growth profile was observed by 10-fold serial dilution of cultures spotted on agar plates with permissive and nonpermissive media, followed by growth at 30°C, 37°C and 16°C temperature (Figure 1(c)). Notably, the growth of the *PGAL7*:Prp16 strain in nonpermissive YPD media is poorer at the low temperature of 16°C as compared to the optimal 30°C. The viability of *PGAL7*:Prp16 strain was significantly reduced as compared to WT only when grown in YPD media and not in permissive YPG media (Figure 1(d)).

**Figure 1.**
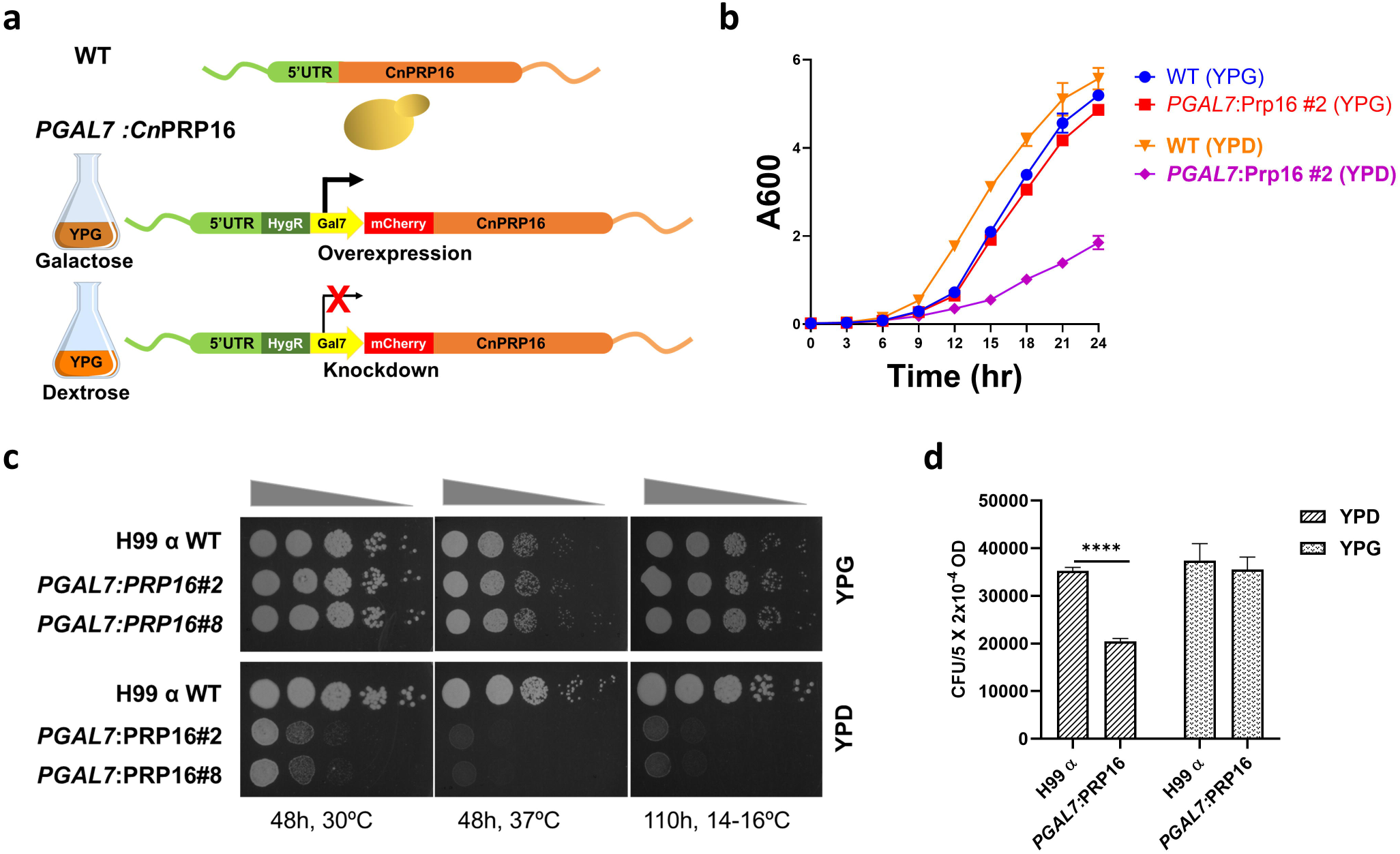
*C. neoformans* Prp16 is essential for growth and viability. (a) Schematic representation of the conditional knockdown *C. neoformans* H99 strain with conditional promoter-shutdown *PGAL7*:mCherry-Prp16 cassette. (b) and (c) The growth kinetics of *C. neoformans* WT and *PGAL7*: Prp16 strains (two independent integrants #2 and #8 with identical growth) in YPG and YPD media. (b) Growth curve at 30°C as recorded by OD_600_ in liquid culture (Error bars represent standard deviation of 3 biological replicates). (c) Growth profile by 10-fold serial dilution on agar plates incubated at 30°C, 37°C and 16°C. (d) Viability of WT and *PGAL7*: Prp16 strains grown in YPG and YPD media for 9 hours and then platted upon YPG plates. Colony forming units (CFUs, shown in y-axis) were calculated from three independent replicate experiments. One-way ANOVA with Tukey’s multiple comparison test was used to calculate significance (ns, P ≥ 0.05; * - P < 0.05; ** - P < 0.01; *** - P < 0.001; **** - P < 0.001).

### Conserved residues in the helicase domains of Prp16 are essential for efficient splicing of cellular introns

Biochemical assays with *S. cerevisiae* Prp16 ascribed its catalytic activity to ATP hydrolysis and conformation remodeling of spliceosome from C complex to C* complex by the unwinding of RNA duplexes [18,19]. The conserved motifs in the Prp16 C-terminal helicase domain are essential for ATP binding, hydrolysis and RNA duplex unwinding. *S. cerevisiae* Prp16 mutants in the conserved motifs cause a dominant negative effect on growth [19]. To gain functional insights into *C. neoformans* Prp16, we generated alanine substitution mutants in its conserved helicase motif 1 (GSGKT) residue K628A and in the motif 2 (DEAH) residue D719A (Supplementary Figure S3(b)). Homologous mutations in *S. cerevisiae* Prp16, K379A and D473A, fail to catalyze the second step of splicing in yeast actin mini-transcripts and competitively inhibit splicing when supplemented into splicing reactions initiated with wild-type yeast extract [18]. In the *PGAL7*:Prp16 strain, we expressed from a heterologous safe haven locus the CnPrp16 wild type, K628A or D719A allele, where their expression was driven by the endogenous Prp16 promoter (Figure 2(a)). These strains with ectopic expression of wild-type or of the mutant CnPrp16 alleles exacerbated the growth defect in *PGAL7*:Prp16 when grown in YPD media (Figure 2(b) and Supplementary Figure S3(c)). The splicing efficiency of selected introns with diverse lengths and varied intronic *cis* consensus elements was tested. The Cln1 intron 5, CNAG_03855 intron 2 and Cas35 intron 7 were chosen to assess their splicing efficiency (Figure 2(c and d) and Supplementary Figure S3(d)). For these introns, accumulation of unprocessed pre-mRNA and a concomitant decrease in spliced mRNA was seen in the *PGAL7*:Prp16 strain grown in YPD media (Figure 2(c and d) and Supplementary Figure S3(e)). Consistent with the rescue of growth defect when wild type Prp16 was expressed ectopically from the safe haven locus, we also note a rescue of the splicing defect of these cellular introns in the *PGAL7*:Prp16 SH:Prp16 strain. Whereas the splicing profile of these introns remained compromised in *PGAL7*:Prp16 SH:Prp16 K628A strain and in the *PGAL7*:Prp16 SH:Prp16 D719A strain (Figure 2(c and d) and Supplementary Figure S3(e)). These data, together with prior literature from *S. cerevisiae* Prp16, suggest that the splicing defects observed in *PGAL7*:Prp16 are dependent on the catalytic activity of CnPrp16.

**Figure 2.**
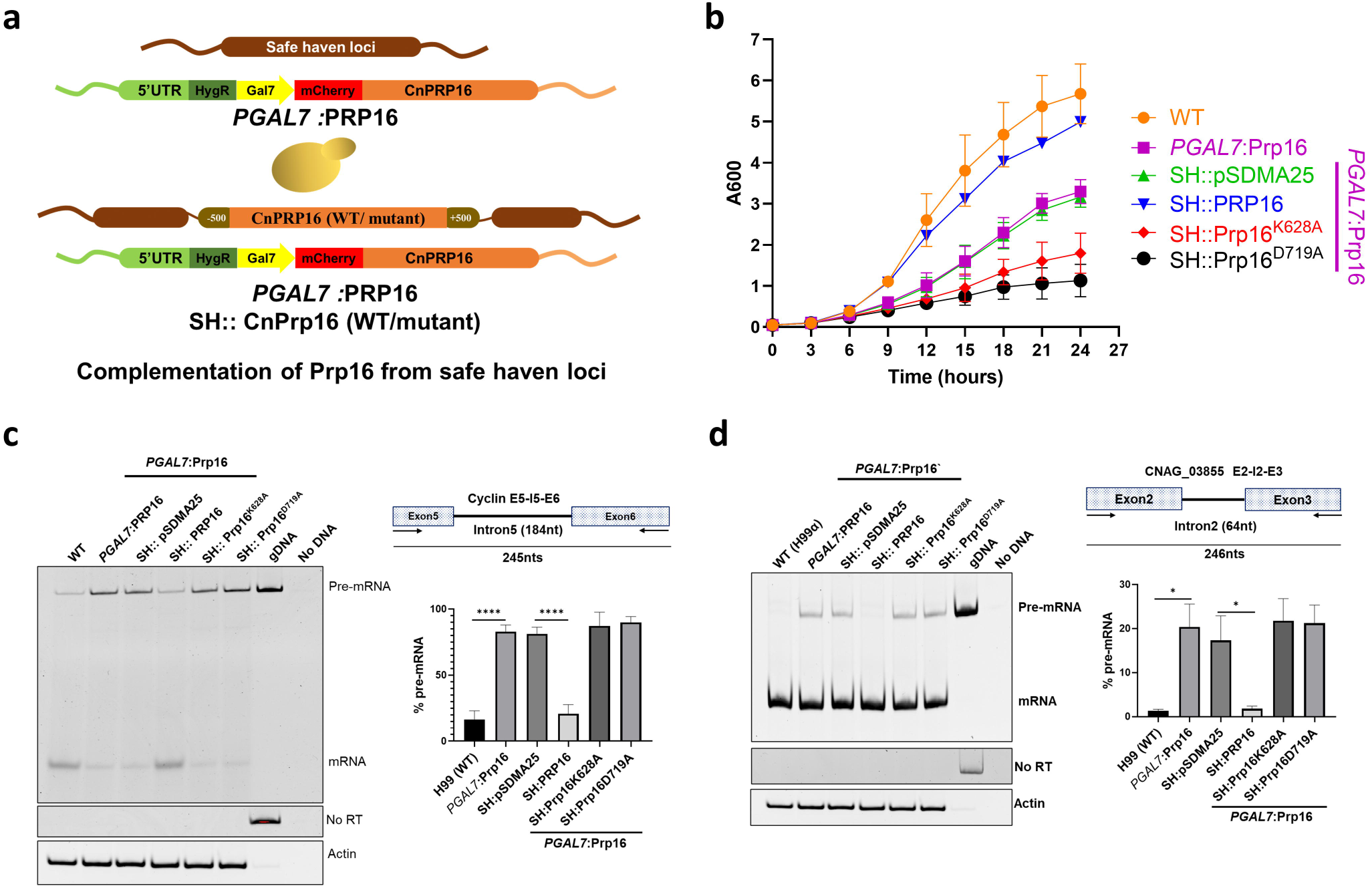
The conserved helicase domain of Prp16 is essential for splicing. (a) Schematic representation of strain expressing full-length CnPRP16 or mutant proteins from the safe haven 1 locus. (b) Growth kinetics of *C. neoformans* WT, *PGAL7*:Prp16 and *PGAL7*:Prp16 strains expressing WT CnPRP16 or its mutants K628A and D719A from safe haven 1 locus. All the strains were grown first in permissive YPG media then subcultured in YPD media for 12 hours. Culture aliquots (at 0.05 OD) were inoculated in YPD media growth measured by recording OD_600_ at 3 hours intervals. The data was plotted with error bars representing standard deviation of 3 biological replicates. (c) and (d) Splicing assays for cellular introns (c) CLN1 Intron5 and (d) CNAG_03855 intron2, by semi-quantitative RT-PCR on RNA isolated from *C. neoformans* WT, *PGAL7*:Prp16 and *PGAL7*:Prp16 strains expressing WT CnPRP16 or the mutants K628A or D719A from safe haven 1 locus. All strains were grown in conditions standardised for Prp16 knockdown (YPD), as described previously. Quantification was done from three biological replicates. One-way ANOVA with Tukey’s multiple comparison test was used to calculate significance (ns, P ≥ 0.05; * - P < 0.05; ** - P < 0.01; *** - P < 0.001; **** - P < 0.001).

### Prp16 knockdown in *C. neoformans* alters genome-wide splicing profile

Next, we assessed the genome-wide splicing consequences of the knockdown of Prp16 in *C. neoformans*. Deep next-generation sequencing of cellular RNAs isolated from three biological replicates of WT and *PGAL7*:Prp16 strains grown in YPD media was done. We achieved high quality and high depth of reads from all samples as is reflected by 60 to 80 million total read counts, of which more than 95% were aligned to the *C. neoformans* genome. As expected, due to promoter shutdown, a decrease in reads aligned to the Prp16 locus was observed for *PGAL7*:Prp16 cells as compared to WT (Supplementary Figure S4(a)). The genome-wide splicing analysis was done by adopting an algorithm previously used for the analysis of the fission yeast *S. pombe* transcriptome [20]. Based on *C. neoformans* genome annotation (GTF) this algorithm uses the numbers of mapped reads traversing exon-exon junctions (EEJR), exon-intron junctions (EIJR) and intron-exon junctions (IEJR) of each intron to calculate the intron retention score (IRS) for each intron. This IRS score is a quantitative representation of splicing efficiency (Figure 3(a)). Introns with the number of total junctions reads < 10 were filtered out from the analysis.

**Figure 3.**
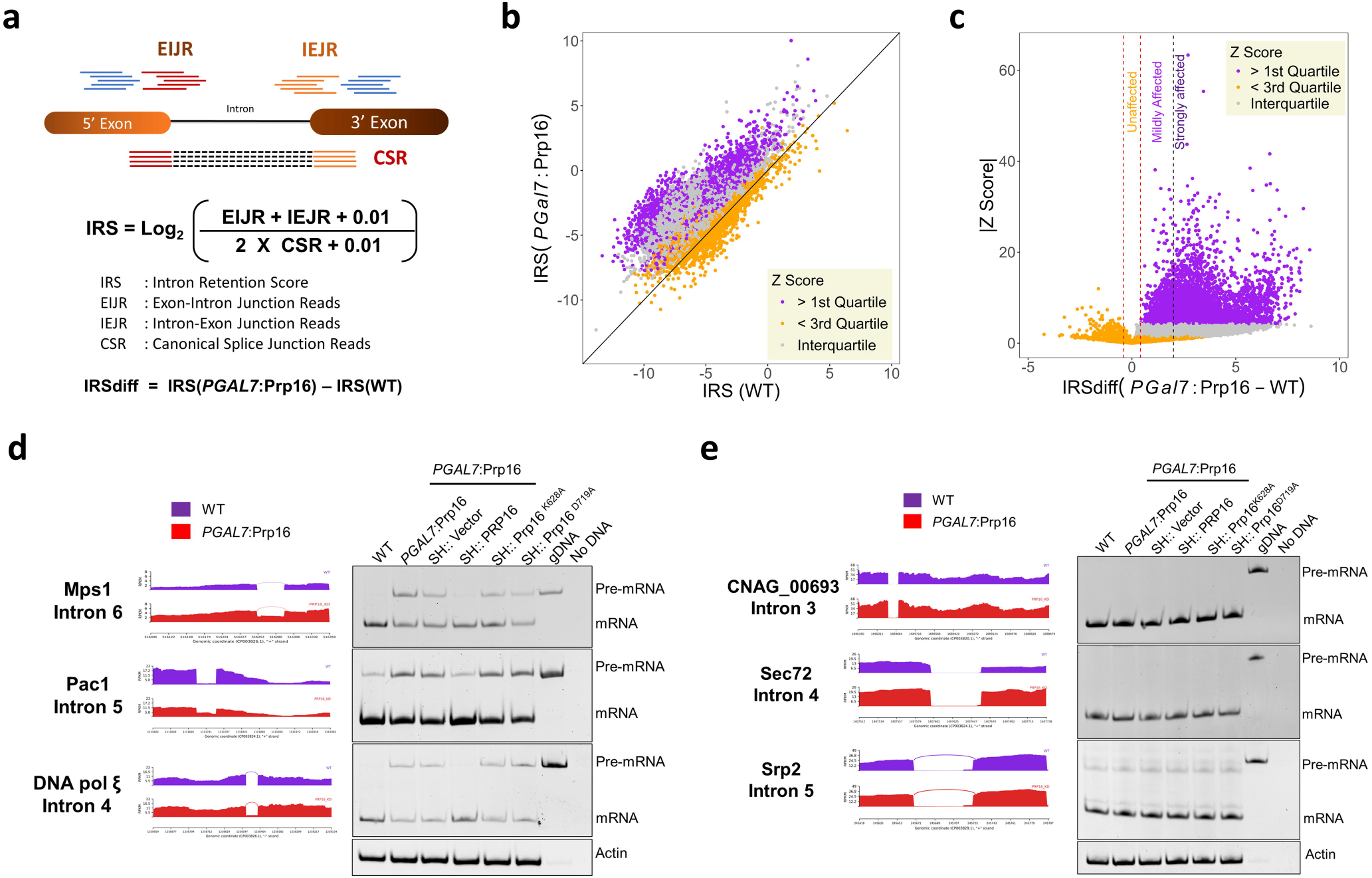
Genome-wide splicing profile on *C. neoformans* Prp16 knockdown. (a) Schematic representation of the algorithm used for genome-wide splicing analysis. (b) Scatter plot of intron retention score (IRS) in *PGAL7*:Prp16 versus WT cells. Each data point represents each annotated intron with a sufficient number of junction reads. The introns are classified based on quartiles of the Z-score of each intron. (c) Plot of absolute Z score against IRSdiff (difference of IRS between *PGAL7*:Prp16 and WT). The vertical red dotted lines represent the cut-off applied to classify unaffected and affected introns, and the vertical dotted black line further demarcates mildly affected and strongly affected introns. (d) Validation of three affected introns with large IRS differences in the RNAseq data and by independent semi-quantitative RT-PCR assays. Splicing defect in the *PGAL7*:Prp16 strain was rescued by ectopic expression of wild type CnPRP16 (SH::PRP16), but not in strains that express helicase domain mutants (SH:Prp16^K628A/D719A^). (e) Validation by semi-quantitative RT-PCR of the splicing status of three unaffected introns in RNAseq data. No pre-mRNA accumulation was seen in the *PGAL7*:Prp16. No pre-mRNA accumulation in strains with ectopic expression of helicase domain mutants of Prp16 (SH:Prp16^K628A/D719A^). For RT-PCR validation, all the strains were grown in conditions standardised for Prp16 knockdown (YPD), as described previously.

An overall shift in the mean IRS values in *PGAL7*:Prp16 cells grown in YPD points to the genome-wide role of Prp16 in the splicing of majority of introns (Figure 3(b)). This reiterates the near universal role for Prp16 for the splicing in *C. neoformans*. As IRS values exhibited a normal distribution (Supplementary Figure S4(b) and S4(c)), two sample Z tests were performed with a null hypothesis of equal IRS between WT and *PGAL7*:Prp16 to calculate the significance of altered splicing and a Z score was assigned to each intron. The values for all introns were sorted by descending intron Z score value and divided into quartiles. Introns with Z score values in > 1st quartile exhibit significant changes in splicing efficiency upon Prp16 knockdown, and introns with Z score values in < 3rd quartile exhibit the least significant changes in splicing upon Prp16 knockdown. This data aligns with the scatter plot of IRS (Figure 3(b)) where introns in > 1st quartile show a greater bias towards Prp16 knockdown (purple data points) and introns in < 3rd quartile lie along the diagonal line (orange data points). The IRS values that fall within interquartile Z score (falling between the 1st and 3rd quartiles) overlaps with the IRS distribution of introns in the > 1st quartile. This may arise from the variations in IRS values observed across different RNA replicates for some affected introns, which result in a lower (interquartile) Z score. For the categorization of affected and unaffected introns, the difference in IRS value in *PGAL7*:Prp16 and WT (IRSdiff) was plotted against the absolute Z score (|Z score|) (Figure 3(c)). Based on this plot, the minimum IRSdiff value obtained for the 1st quartile was assigned as a cutoff between affected and unaffected introns (Figure 3(c), vertical red dotted lines). Affected introns were further divided into strongly affected and mildly affected introns based on whether the IRSdiff value was greater than or less than 2, respectively. This analysis reveals that > 88 percent of introns (25,642) were affected by the knockdown of Prp16, and < 12 percent (2808) remained unaffected upon Prp16 knockdown (Supplementary Figure S4(d)). A small set of 685 introns had a negative IRSdiff value and a negative Z score. This group we surmise could be false positives from the unaffected category of introns but where transcripts had either an increase in mRNA read count or a decrease in pre-mRNA read count. This set of 685 introns were not taken for further analysis.

The data from the bioinformatic analysis was validated by semi-quantitative RT-PCR for some introns chosen from the strongly affected and the unaffected categories (Figure 3(d) and 3(e)). Consistent with the transcriptomic data Mps1I6 (Mps1 Intron6), Pac1I5 (Pac1 Intron5) and DNApolξ I4 (DNApolξ Intron 4) from strongly affected category showed RT-PCR amplicons denoting increased pre-mRNA accumulation in *PGAL7*:Prp16 cells grown in YPD media. Expression of wild-type Prp16 from the heterologous safe haven locus rescued the impaired splicing of all three introns (Figure 3(d), lane 4). Ectopic expression of helicase DEAD box mutant alleles: Prp16 K628A and D719A, from safe haven locus (*PGAL7*:Prp16 SH:Prp16 K628A strain and *PGAL7*:Prp16 SH:Prp16 D719A) could not rescue the splicing defect triggered by the knockdown of Prp16 as evident from high levels of unspliced precursor (Figure 3(d), lane 5 and 6). For the introns CNAG_00693I3, Srp2I5 and Sec72I4 from the unaffected category, RT-PCR assay also showed the splicing profile was unaltered in *PGAL7*:Prp16 cells grown in repressive media (Figure 3(e)). Interestingly, splicing of these introns remained unaffected even upon ectopic expression helicase DEAD box mutant alleles from the safe haven locus in *PGAL7*:Prp16 cells. This gives the hint that splicing of these introns could be Prp16-independent splicing, as the predicted dominant negative effect of K628A and D719A mutants was not observed.

### Exonic consensus elements at pre-mRNA 5’ splice site and 3’ splice site corelate with Prp16 dependent efficient splicing

Next, we assessed the *cis* features in introns, and at exon-intron junctions, for introns where splicing was strongly compromised upon Prp16 knockdown as compared to introns where splicing was unaffected. To this end we characterized the 5’ splice site (5’SS) and 3’ splice site (3’SS) of all ~40,000 *C. neoformans* introns, 4954 introns strongly affected introns (Z score > 1^st^ quartile) in *PGal7*:Prp16 cells and the set of 2808 unaffected introns (Figure 3(c) Supplementary Figure S4(d)). As per *C. neoformans* genome annotation, sequence logo was generated for 5’SS and 3’SS of all three intron groups (Figure 4(a) and 4(b)). Then differential enrichment or depletion of specific nucleotides around the 5’SS and 3’SS of strongly affected and unaffected introns was visualized using DiffLogo (Figure 4(c)). Notably, the last 3 nucleotides preceding the 5’SS are enriched for AAG in the set of unaffected introns and the 1^st^ nucleotide of 3’exon *i.e*., downstream of 3’SS more frequently a G in the unaffected introns (Figure 4(c)). Further, we deduced that intronic N3 and N4 nucleotides at the 5’SS being ‘GA’ is also a signature common to affected introns (Figure 4(c)). No differential enrichment was observed between affected and unaffected introns for intronic sequences at the poly-pyrimidine track and 3’SS preceding the 3’ exon (Figure 4(c)).

**Figure 4.**
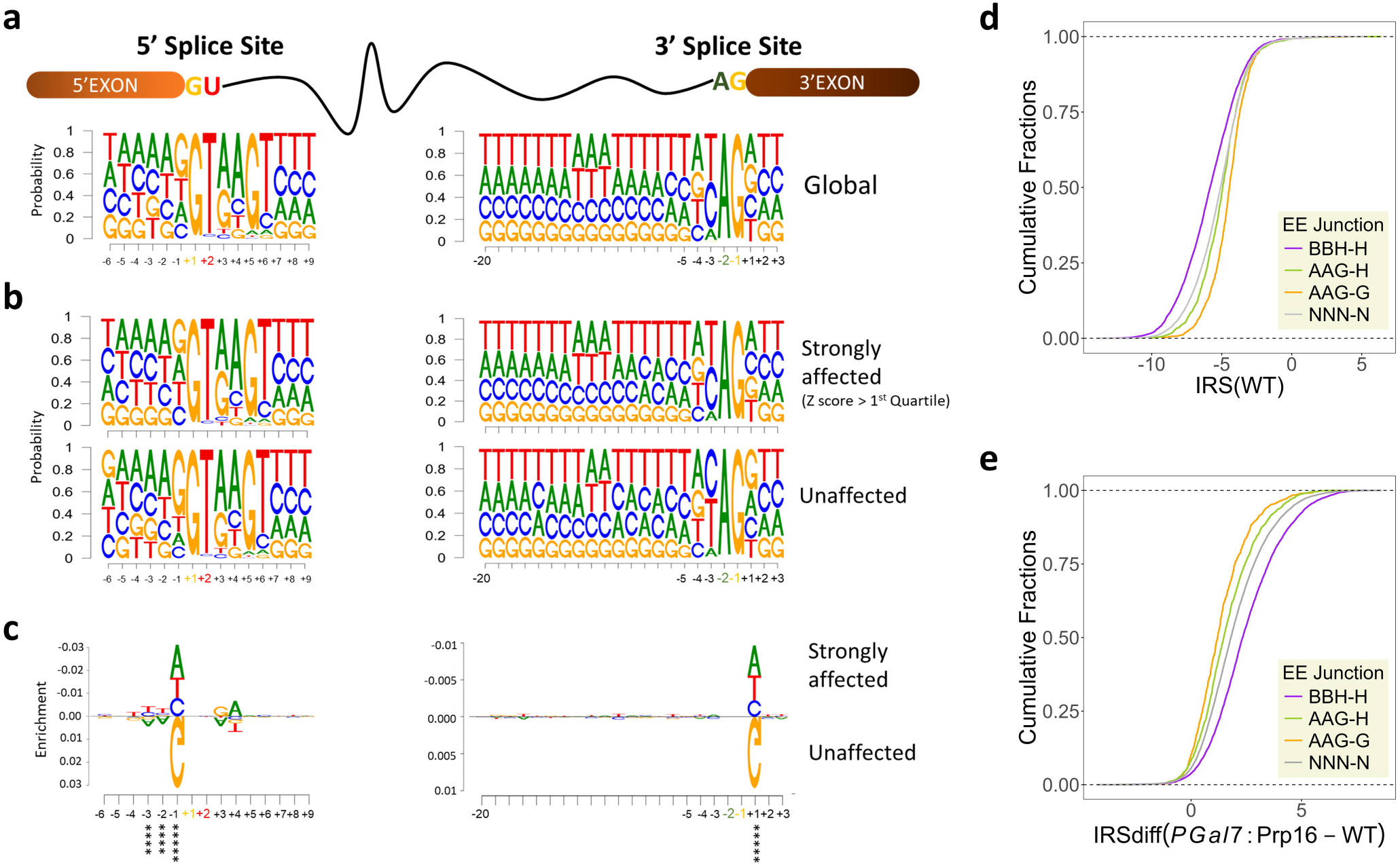
Intronic features determining intron dependency on Prp16. (a) Sequence logo for 5’ splice site and 3’ Splice site for total annotated *C. neoformans* introns. (b) Sequence logo for 5’ splice site and 3’ Splice site for strongly affected introns with Z score >1^st^ quartile (upper panels) and unaffected introns (lower panels). (c) DiffLogo analysis was comparing differential sequence enrichment of the 5′SS and 3’SS between strongly affected introns high Z score (>1^st^ quartile) and unaffected introns. The enrichment of A, A and G against other nucleotides at positions −3, −2, and −1 of 5’SS of unaffected introns was significant by p-values under 6.9C×C10^−12^, 7.2C×C10^−14^ and 3.9C×C10^-105^ by two-sided Fisher’s exact test, respectively. The enrichment of A against other nucleotides at positions +1 of 3’SS of unaffected introns was significant by p-values under 1.5C×C10^-27^ by two-sided Fisher’s exact test. (d) Cumulative plot of IRS in WT for each group of classified introns. All introns are classified into four groups based on their last three nucleotides of 5’exon and 1^st^ nucleotide of 3’exon. (e) Cumulative plot of IRS differences for each group of classified introns in *PGAL7*:Prp16 versus WT.

*In vitro* splicing assays for Prp16 activity using human or *S. cerevisiae* splicing extracts showed its role for remodeling splicing complexes that allows 5’ exon and 3’ exon in a conformational alignment that promotes the second step of catalysis. This is facilitated by U5-snRNA loop1 interactions with the 5’ exon and 3’exon [21]. Therefore, we focused our next analysis on *cis* sequences located at the end of 5’ exons and at the start of 3’ exons, to decipher any differentially enriched in unaffected introns as compared to strongly affected introns. We classified introns into four groups based on the last three nucleotides of the 5’ exon and the first nucleotide of the 3’ exon as: AAG-G (872 introns), AAG-H (1896 introns), BBH-H (5596 introns), and NNN-N (20671 introns) respectively. Next, we assessed the splicing efficiency of these intron classes by calculating IRS in WT and plotting cumulative distribution plots. Notably, in the WT strain, introns with the BBH-H signature are the most efficiently spliced, while those with the AAG-G signature are the least efficiently spliced introns (Figure 4(d) and Supplementary Figure S5(a)). The splicing defect upon Prp16 knockdown was assessed for these classes by calculating IRSdiff and plotting cumulative distribution plots. Consistent with the consensus sequence seen by seqLogo and DiffLogo algorithms, we see introns with AAG-G signature are the least affected and the BBH-H signature are the most affected on depletion of Prp16 (Figure 4(e) and Supplementary Figure S5(b)). This suggests Prp16 has a role in enhancing the efficiency of splicing for introns with sequence diversity in the last three nucleotides of the 5’ exon that precede the 5’SS and the first nucleotide of 3’ exon that follows the 3’SS.

### Pre-mRNA substrates with weaker interactions with U5 snRNA loop1 are Prp16-dependent

To experimentally validate the interpretations emerging from the bioinformatic analysis of splice sites in Prp16 dependent and independent introns, we designed minigenes to be tested for their splicing efficiency in *C. neoformans* cells. We chose Pac1 intron5 and its flanking exons (E5I5E6) from the set of strongly affected introns in the Prp16 knockdown dataset. The construct containing this minigene under control of the H3 promoter was cloned in an integration vector pEE27 with flanking sequences suitable for integration to heterologous safe haven 3 locus [22]. Thus, once integrated into WT or *PGal7*:Prp16 strains, mini-transcript expression and its splicing status can be assessed. We find the mini-transcript splicing efficiency parallels that seen for the intron 5 in the endogenous Pac1 transcript (Supplementary Figure S5(c) and S5(d)). In these splicing mini-transcript assays, we also generated and tested two variants. The first where the last three nucleotides of 5’ exon in the mini-transcript were mutated from UUC to AAG and the second where these 5’exon mutations were combined with A to G change in the first nucleotide of 3’ exon of the mini-transcript (Figure 5(a)). These variant mini-transcripts were also expressed from the safe haven 3 locus in WT and in *PGal7*:Prp16 cells. The strong splicing defect of the mini-transcript in Prp16 depleted cells is alleviated when the 5’SS exonic UUC sequence was mutated to AAG, as the mini-transcript intron was now spliced more efficiently in *PGal7*:Prp16 cells grown in YPD (Figure 5(b), lane 2 *vs.* lane 4, Bar graphs pink *vs.* green). For mini-transcripts with A to G mutation in the 1^st^ nucleotide of 3’exon that also had 5’SS exonic UUC to AAG nucleotide mutations (Figure 5(b), lane 6), the splicing efficiency was marginally improved as compared to mini-transcript with only 5’exonic mutations (compare *PGal7*:Prp16 lanes 4 *vs.* 6 Bar graphs green *vs.* orange). However, this improved splicing efficiency of the AAG-G mutant Pac1 I5 mini-transcript under conditions of Prp16 depletion did not approach the efficiency of splicing seen for other endogenous cellular transcripts with the AAG-G *cis* signature, which are spliced well even after Prp16 depletion. This suggests that additional factors apart from exonic sequences at 5’SS and at 3’SS could contribute to the requirement of Prp16 for efficient splicing.

**Figure 5.**
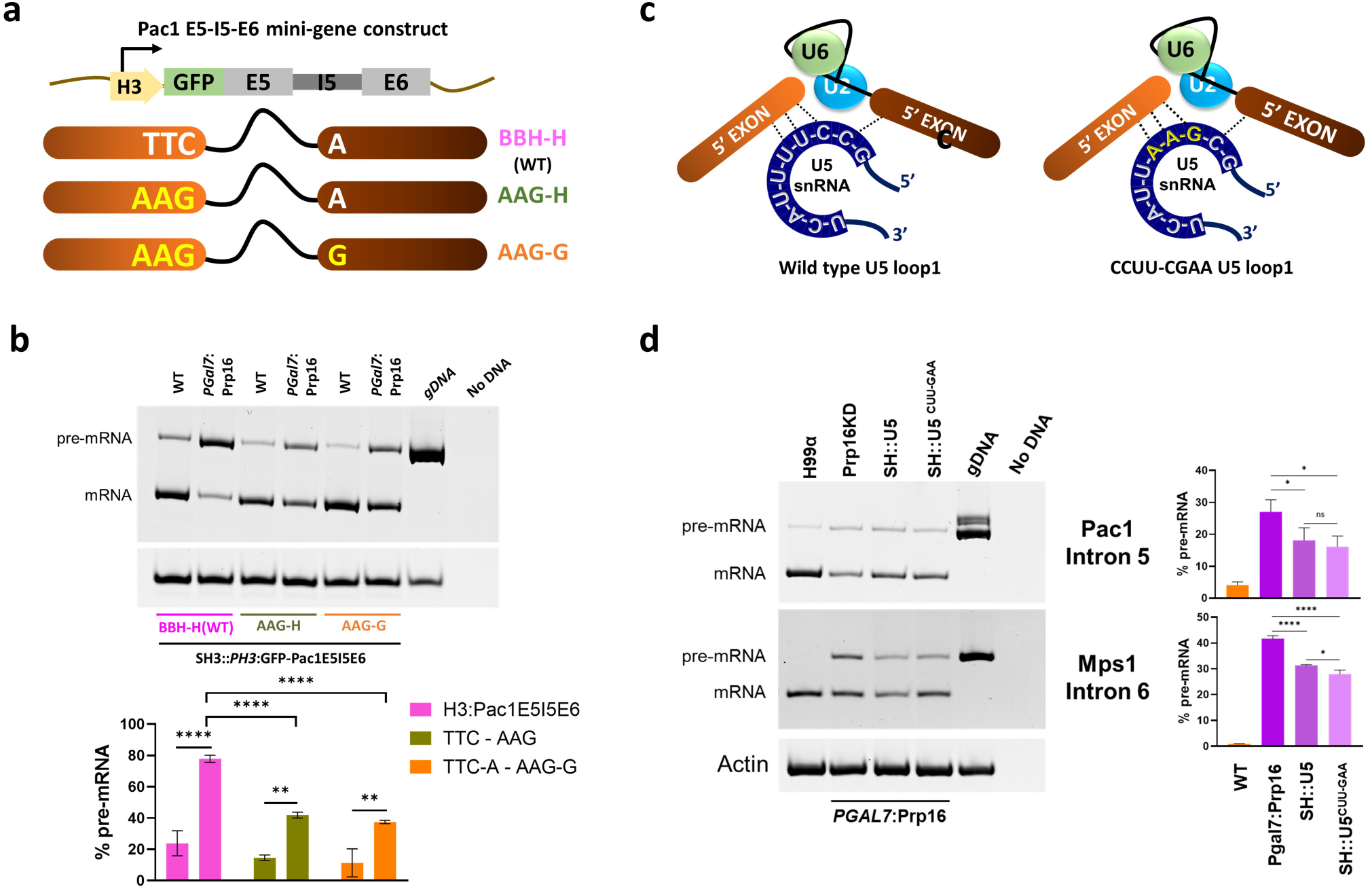
Interactions between U5 loop1 and pre-mRNA contribute to Prp16 dependency (a) Schematic representing the Pac1E5I5E6 minigene construct and mutants generated at 5’ exon and 3’exon. (b) Semi-quantitative RT-PCR for Pac1E5I5E6 minigene expressed from safe haven locus in WT and *PGAL7*:Prp16 strains. (c) Schematics for U5 pre-mRNA interactions before exon ligation for wild type U5 and loop1 mutant U5^CUU-GAA^. (d) Semi-quantitative RT-PCR of cellular Pac1E5I5E6, Mps1E6I6E7 in WT, *PGAL7*:Prp16 and *PGAL7*:Prp16 strains ectopically expressing wild type U5 and U5^CUU-GAA^ from safe haven locus. Semi-quantitative RT-PCR was quantified from three biological replicates. One-way ANOVA with Tukey’s multiple comparison test was used to calculate significance (ns, P ≥ 0.05; * - P < 0.05; ** - P < 0.01; *** - P < 0.001; **** - P < 0.001).

As an orthogonal approach, we examined whether U5 snRNA interactions with pre-mRNA *cis-*elements could influence splicing status of introns that are not spliced well under conditions of Prp16 knockdown. The U5 locus, along with the 500 bp upstream genomic region (as an endogenous promoter) and 500 bp downstream genomic region was cloned into the integration vector pSDMA57 that is specific for integration into the safe haven 1 locus [23]. We also created mutants in U5 loop 1 residues in the pSDMA57 clone that likely improve base-pairing interaction with exonic sequences in cellular transcript for Pac1E5I5E6 pre-mRNA (Supplementary Figure S6(b)). To achieve this we mutated cloned U5 sequence in pSDMA57 so as to express mutant U5 with loop1 sequence as GAA (Figure 5(c)). The U5^WT^ and U5^CUU-GAA^ mutant expression constructs in the pSDMA57 vector were integrated at the safe haven 1 locus of *PGAL7*:Prp16 strains (Supplementary Figure S6(a)). In RT-PCR assays, we observed that the expression of this additional copy of wild type U5 and U5^CUU-GAA^ in *PGAL7*:Prp16 strains improved the splicing efficiency of cellular Pac1 E5I5IE6. This improved splicing is manifested by increased mRNA, but no change in pre-mRNA was observed (Figure 5(d) lane 2 *vs.* lane 3). This may point to a rescue/ improved efficiency of the second step of splicing for Pac1E5I5E6. Consistently, in *PGAL7*:Prp16 cells with the expression of U5^CUU-GAA^ mutant, we note slightly better splicing of cellular Pac1 E5I5IE6 as compared to *PGAL7*:Prp16 cells that express wild type U5 from safe haven locus (Figure 5(d), top panel lane 2 *vs.* lane 4). We also examined the splicing of Mps1 E6I6E7, where pre-mRNA interactions with U5 loop 1 are not canonical Watson Crick base pairing interactions. The splicing efficiency was assessed in cells expressing either U5 or U5^CUU-GAA^ from the safe haven locus (Supplementary Figure S6(b)). As seen for Pac1 I5 splicing, we note improved splicing of Mps1 E6I6E7 upon expression of the additional copy of wild-type U5 in *PGAL7*:Prp16 cells (Figure 5(d), lower panel lane 2 *vs*. 3). Further, in *PGAL7*:Prp16 SH:U5^CUU-GAA^ cells where U5 loop1 interactions with Mps1 E6I6E7 are altered we note a marginally better splicing of Mps1 intron 6 as compared to its splicing in *PGAL7*:Prp16 SH:U5 cells (Figure 5(d), lane 3 *vs.* 4). It is plausible that U5^CUU-GAA^ transcript levels, as expressed from U5 integrated mini-gene are inadequate to efficiently compete with endogenous U5 and consequently to have only subtle changes to the splicing status of cellular transcripts. These results support the hypothesis that stronger pre-mRNA recognition by increased expression of wild-type U5 snRNA, or even the U5^CUU-GAA^ mutant, may suppress intron retention caused by Prp16 depletion.

### Prp16 promotes the splicing of short and ultra-short introns in *C. neoformans* genome

The EM structure of human spliceosomal complex A formed after assembly of U1 and U2 snRNPs covers approximately 79-125 nts of RNA substrate [24]. The remarkable variability of intron lengths from < 50 nts to > 50,000 nts [25] seen in several genomes poses questions on the assembly of the spliceosomal complex on short and ultra-short introns without stearic hindrance. Comparative analysis of intron length distribution across diverse species revealed a higher prevalence of short and ultra-short introns in most fungal genomes studied thus far. Strikingly, *C. neoformans* harbors the highest percentage of short (60-100 nts) and ultra-short (<60 nts) introns among all the studied fungal species (Figure 6(a)). In the dataset of introns affected by CnPrp16 depletion, we observed a higher representation of short introns in the strongly affected and mildly affected introns categories than in unaffected group of introns (Figure 6(b)). We analyzed the splicing efficiency of introns after categorizing them based on intron length using finer bin sizes for the ultra-short intron group. The CDF plot of IRS values in the WT dataset (Figure 6(c)) suggests *C. neoformans* ultra-short introns 40-60nt (dark red line) are spliced most efficiently, followed by short introns 60-100nt (light green line), introns in range 100-500 nts (orange line), and the poorest efficiency even in wild type cells is for long introns > 500nt (grey line). A small population of introns shorter than 40nt (purple lines) shows greatest IRS value, suggesting this group to have notably poor splicing efficiency. This data opens an intriguing interpretation that *C. neoformans* spliceosomal machinery, being apparently geared for recognizing and splicing a majority of its genome’s ultra-short and short introns (covering lengths of 40-100nts). Next, we also analyzed splicing defects in introns binned into groups based on their length in Prp16 knockdown conditions. Here we calculated and plotted the IRSdiff value for introns categorized based on size (Figure 6(d)), and we observed a higher splicing defect (IRSdiff) for ultrashort introns (20-40 nts and 40-60 nts, purple and dark red lines) followed by short introns (60-100 nts, light green line). This correlation between splicing defect upon depletion of Prp16 (IRSdiff) and intron length is significant when examined even scanned at 10 nts resolution (Supplementary Figure 6(c) and 6(d)). This correlation was seen only for introns from 20nt to 80nt introns as no significant differences in IRSdiff scores were observed for introns classes 80-90 nts, 90-100 nts and those > 100 nts (Figure 6(d) right panel and Supplementary Figure 6(d)). Thus a role of Prp16 in the efficient splicing of ultra-short and short introns in *C. neoformans* transcriptome is deduced.

**Figure 6.**
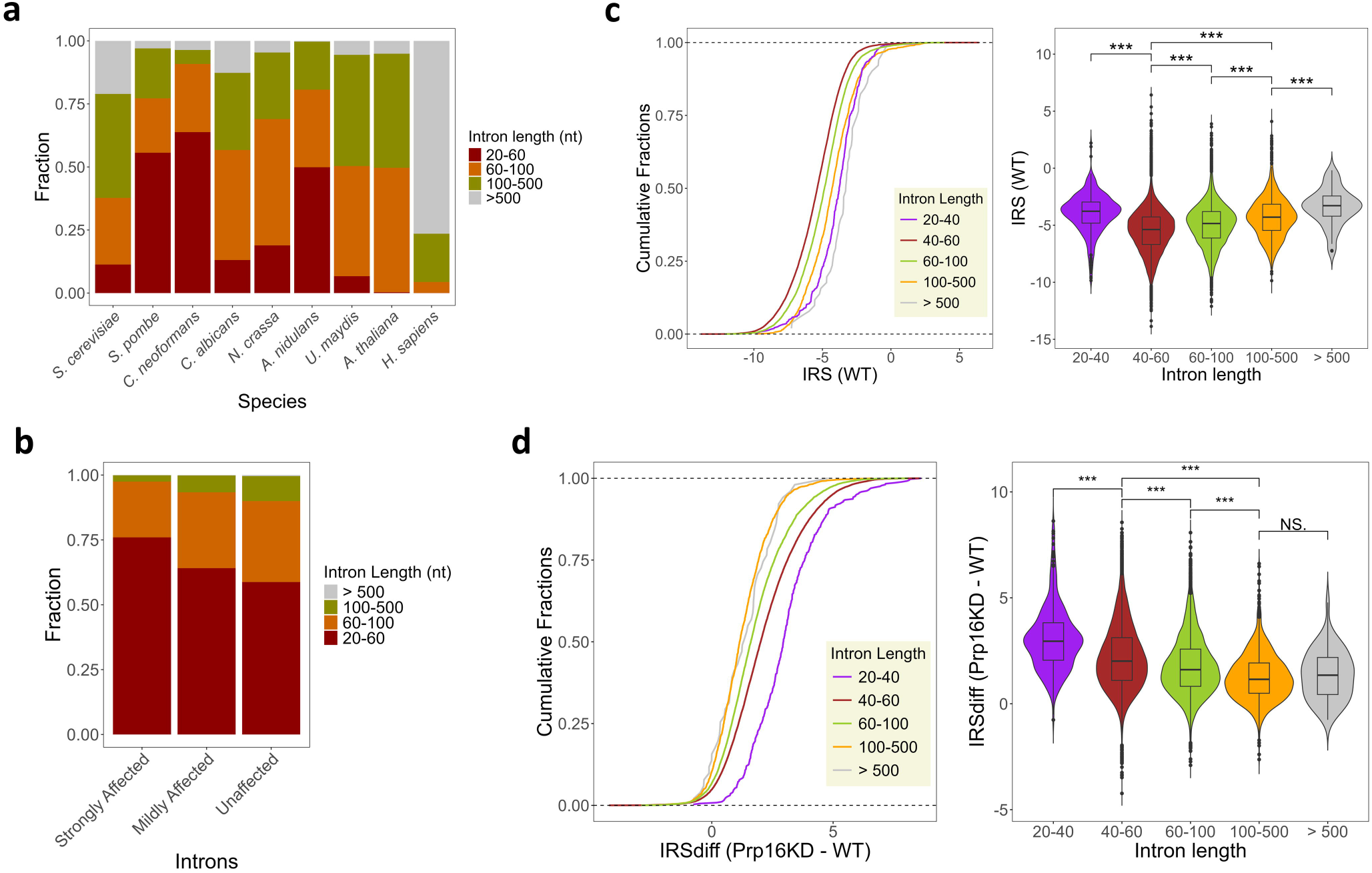
(a) Stacked bar plot for intron sizes across different species. *C. neoformans* possess a maximum fraction of short and ultra-short introns among all species. (b) Stacked bar plot for the occurrence of various size introns in the transcriptome of Prp16 knockdown classified as strongly affected, mildly affected and unaffected. (c) Cumulative plot of intron retention score (IRS) for each group of classified introns in WT transcriptome. All introns are classified into four groups based on their length: 20-40 nts, 40-60 nts, 60-80 nts, 80-100 nts, 100-500 nts and >500 nts. The right panel shows a box plot of IRS in WT for each group of introns. (d) Cumulative plot of IRS difference between *PGAL7*:Prp16 and WT for each group of classified introns. The right panel shows a box plot of IRS differences between *PGAL7*:Prp16 and WT for each group of introns. Asterisks indicate statistically significant differences, as determined by the Wilcoxon rank-sum test within the R package ggplot2.

## Discussion

The dynamic assembly, activation and catalysis of the spliceosome involves several remodeling steps driven by eight helicases, including DEAD and DEAH-box helicase proteins [4]. Prp16, one of the DEAH-box helicase, is detected in step one spliceosomes by the cryo-EM of human and yeast complexes [26,27]. Single-molecule and cross-linking experiments suggest that Prp16 binds single-stranded RNA region downstream of the branchpoint and translocate on the lariat intermediate as a molecular winch [28,29]. The ATPase activity of Prp16 facilitates the conversion of the branching C complex to the step two catalytically activated C* complex [30]. The role of splicing factors in development and diseases are being reported from several models [31–33]. Interestingly, the autosomal recessive mutation G332D in DHX38, the human orthologue of Prp16 is associated with retinitis pigmentosa [34], yet its requirement for introns with diverse sizes and splicing signatures remains understudied.

In this study, we report the genome-wide splicing defects in the short intron-rich genome of *C. neoformans* that are triggered by Prp16 knockdown. We find that a small subset of introns with flanking exonic sequences that can be robustly recognized by U5 snRNA are spliced well in Prp16 knockdown conditions. While complementation of splicing defects in knockdown conditions is achieved by the expression of wild-type Prp16 from a heterologous locus, *C. neoformans* strains expressing Prp16 mutants in the catalytic motif I (K628A) and motif II (D719A) grow poorly and remain splicing defective. Interestingly, these phenotypes are similar to observations on the *S. cerevisiae* Prp16 mutants K379A and D473A, in homologous residues, which are synthetic lethal in the *S. cerevisiae prp16*Δ null mutant and cause dominant negative growth impairment when overexpressed in wild-type yeast [19]. *In vitro,* these *S. cerevisiae* Prp16 mutant proteins are unable to catalyze the second step of splicing in budding yeast splicing extracts depleted of Prp16 and further when supplemented to reactions with wild-type extracts these mutant proteins inhibit splicing [18]. Consistent with these observations, K628A and D719A alleles of *C. neoformans* Prp16 exacerbate the growth defect of *PGAL7*:Prp16 strains grown in nonpermissive media, indicating dominant negative growth suppression as observed in *S. cerevisiae.* Further, K628A and D719A alleles of *C. neoformans* Prp16 do not alter the splicing defects triggered by Prp16 knockdown cells, suggesting that the splicing defects due to Prp16 depletion are predominantly associated with the catalytic activity of *C. neoformans* Prp16.

Genome-wide transcriptome analysis revealed that Prp16 is required for splicing of the vast majority of *C. neoformans* introns. Interestingly, a small subset of introns is spliced efficiently even after Prp16 depletion, and these introns are also spliced in strains that express Prp16 K628A and D719A mutants. A significant proportion of these introns spliced independent of Prp16 are enriched for AAG residues as the last three nucleotides of their 5’ exon (−3, −2 and −1 positions of 5’SS) and with G residue as 1^st^ nucleotide of their 3’ exon (+1 position following the 3’SS). In *S. pombe* and *A. thaliana,* introns where the 5’exons end with AAG are strongly recognized by U5 snRNA and can suppress the splicing defect triggered by mutations in U6 m6A methyltransferase [20,35]. Statistical analysis of human transcriptome suggests that G nucleotide at the end of 5’ exon and at the start of 3’ exon favors a strong binding register for U5 loop1 C39|C38 and significantly contributes to specific exon recognition and splicing precision [21]. Our analysis reveals that U5 snRNA-pre-mRNA intron interaction in *C. neoformans* transcripts is one factor determining the requirement of Prp16 for efficient splicing of each intron. In budding yeast *in vitro* splicing assays the U5.U6/U4 tri-snRNP assembled in the spliceosome has confirmation that allows U5 loop 1 recognition of these terminal three nucleotides of the pre-mRNA 5′ exon. After first step of catalysis, this interaction ensures the 5’ exon is tethered to the spliceosome. This interaction also aligns and allows U5 interaction with 3’ exon in the C* complex for second-step catalysis [36,37]. Our global transcriptome study inferred that pre-mRNA exonic AAG-G sequences that have a strong interaction with U5 loop 1 could favor conformational transitions even when *C. neoformans* cells are depleted for Prp16 activity. On the contrary, introns with a weaker pre-mRNA exonic interactions with U5 loop1 (*e.g.,* BBH-H) require Prp16-driven remodeling for spliceosomal transitions (Figure 7). This inference is consistent with other data from our mutational analysis of mini-transcripts where “BBH-H” exonic signatures were mutated to “AAG-G” in a model minigene and were tested in cells depleted of Prp16. The significantly enhanced splicing efficiency of the mutant mini-transcript supports the interpretation that strong U5 loop 1 interactions with 5’ exon could strengthen the anchoring of 5’SS for spliceosome activation. Since we note the conversion of only 5’ exon “BBH” to “AAG” signatures also enhances the splicing of mini-transcript in Prp16 knockdown conditions this opens the speculation of a role for *C. neoformans* Prp16 prior to first step splicing catalysis. We also report that increased U5 snRNA levels can improve the splicing efficiency of Prp16-dependent introns. These data and our studies on the effect of U5 loop 1 mutants supports the hypothesis that the interaction of U5 snRNA with pre-mRNA contributes toward the requirement of Prp16 for efficient splicing in *C. neoformans* transcripts.

**Figure 7.**
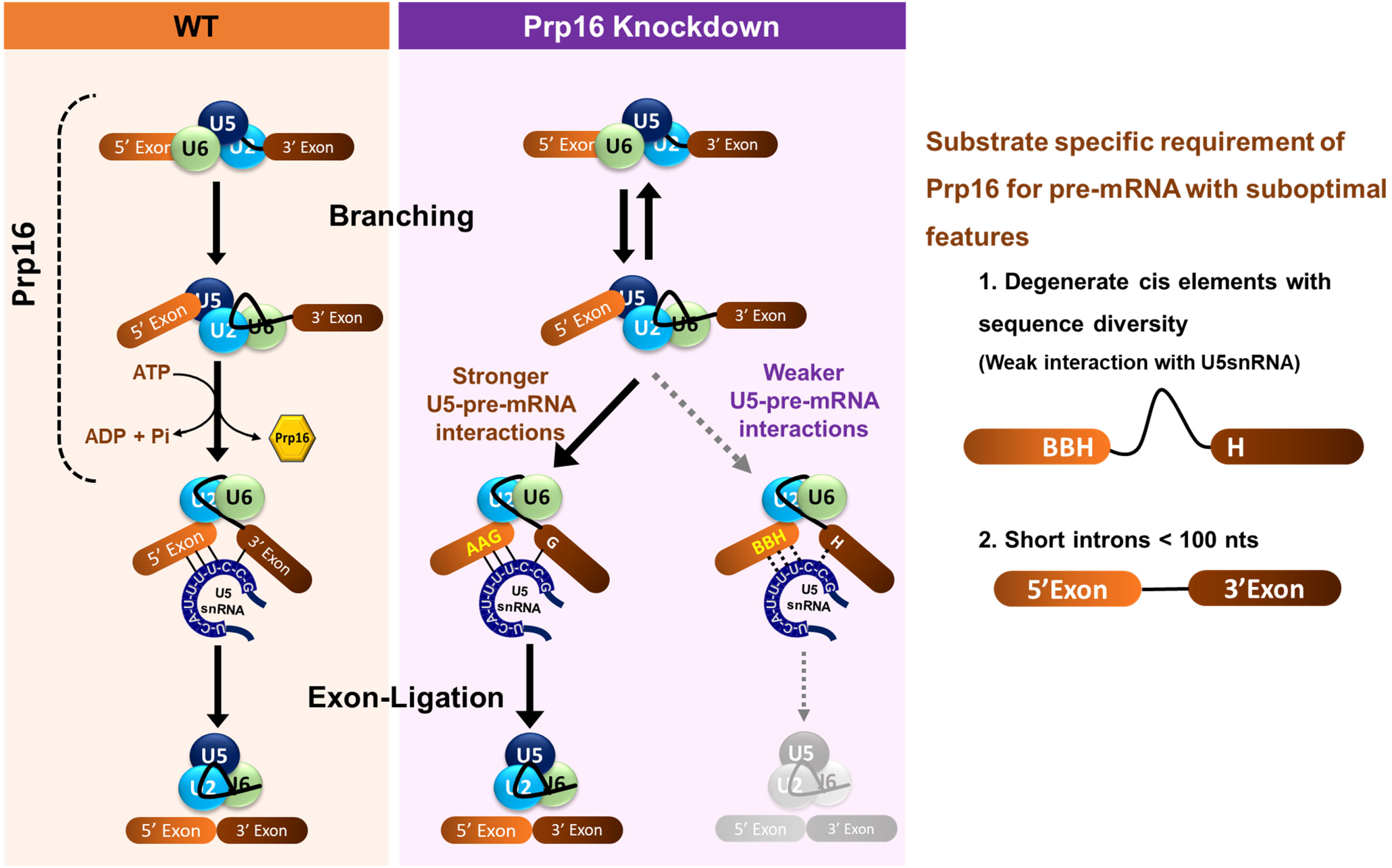
Schematic illustration of the two spliceosomal catalytic reactions in WT and Prp16 knockdown condition. In WT, ATP-dependent helicase activity of Prp16 catalyzes conformational transition from branching conformation after first step catalysis to exon ligation conformation primed for the second step. In Prp16 knockdown conditions (*PGAL7*:Prp16), these conformational transitions are efficient for introns with stronger U5 loop1-pre-mRNA interactions (AAG-G). However, introns with weaker U5 loop1-pre-mRNA interactions (BBH-H) require Prp16 to remodel the branching conformation to exon ligation conformation. The right panel depicts that Prp16 enables the splicing of introns with diverse exonic sequences at 5’SS and 3’SS. Additionally, Prp16 also requires the splicing of short and ultra-short introns.

A notable feature of the *C. neoformans* transcriptome is the high prevalence of short introns. In the *C. neoformans* genome, the average intron size is 56 nts [38]. The human spliceosomal complex A with U1 and U2 snRNPs is a ~26C×C20 × 19.5 nm globular and asymmetric particle that can cover 9-125 nts of linear RNA upon assembly [24,39]. However, all organisms possess some fraction of introns in their transcriptome that are much shorter than 79 nts. For instance, studies have reported human ultra-short introns with lengths as short as 43–65 nts, that are regardless efficiently spliced [25,40]. The underlying mechanisms for recognizing the *cis*-elements in the short introns by the spliceosome without steric hindrance remain unknown. A recent study uncovered that the human alternative splicing regulator SPF45 (RBM17) is essential for efficiently splicing many short introns. SPF45 competes with U2AF^65^ for binding to the U2 snRNP protein SF3B1 and facilitates spliceosomal assembly on short introns with short or truncated polypyrimidine tracks [24]. The ultra-short intron-rich genome of *C. neoformans* is equipped with a spliceosomal machinery that is efficient for recognition and splicing these ultra-short introns. As we observe in our study, that there is an increased proportion of short and ultra-short introns among the group of affected category of introns on depletion of Prp16, the data suggests a role of Prp16 in splicing *C. neoformans* ultra-short introns. In budding yeast, spliceosomal helicases like Prp16 and Prp22 bind to spliceosome transiently and at a distance downstream from the RNA structure that they remodel [28,29]. This feature eliminates the intron size restriction for Prp16-mediated spliceosome remodeling and opens avenues for future studies in *C. neoformans* to understand the mechanism of splicing of ultra-short introns that could rely on *in vivo* approaches on model mini-transcripts.

Taken together, our study uncovers the genome-wide but substrate-specific function of Prp16 in *C. neoformans*. Intronic features associated with the strength of U5 loop 1–pre-mRNA interactions determine the requirement of Prp16 for splicing and are likely relevant for spliceosomal conformational transitions to ensure splicing efficiency. Further, we underscore the role of Prp16 in splicing ultra-short introns prevalent in the *C. neoformans* genome. This work also shows that pre-mRNA substrates possessing optimal intronic features, such as a strong snRNA binding register and sufficient length, are capable of undergoing splicing even in a minimal spliceosome. Conversely, sub-optimal introns necessitate the involvement of splicing factors, such as Prp16, for stable non-Watson-Crick interactions with the spliceosomal snRNAs for efficient splicing.

## Materials and Methods

### Yeast strains, primers, and media used in this study

The strains and primers used in this study are listed in Supplementary Table S1 and Supplementary Table S2, respectively.

### Construction of *C. neoformans* Prp16 promoter shutdown conditional knockdown

To generate promoter shutdown conditional knockdown of Prp16, we replaced the endogenous promoter of Prp16 with the GAL7 promoter. The 1KB fragment upstream of the Prp16 start codon was PCR amplified from the H99 genomic DNA and cloned into the pBSKS+ vector using the oligos listed in S2 table. Similarly, 1KB fragment from the start codon of Prp16 was amplified by PCR and cloned into the pBSKS+ vector. After validation by sequencing, the two fragments were subcloned into the pGAL7 mCherry vector with a hygromycin selection marker. The upstream fragment was cloned at the SacI site, and the downstream fragment was cloned at the HindIII-XhoI site to obtain the final clone where the reading frame of mCherry is in translational fusion with the N-terminal of Prp16. The resulting plasmid was partially digested with SacI, and the other SacI at the 3’ end of the upstream fragment was deleted by end filling. The resulting construct was digested to release the homologous recombination cassette was released with SacI and Xho1 digestion and introduced into wild-type H99 cells by biolistic transformation. The hygromycin-positive transformants were confirmed locus-specific PCR using the oligos listed in Supplementary Table S2.

### Construction of safe haven expression constructs for complementation or for expression of mini-transcripts

To complement the Prp16 knockdown strain with wild type Prp16, the CnPRP16 locus was cloned along with 500bp upstream and 500bp downstream intergenic region in safe haven integration vector pSDMA25. To generate mutants of Prp16 at specific amino acids in the conserved catalytic motifs, overlap PCR was done using mutagenic primers specific to K628A and D719A sites along with Prp16 primers listed in Supplementary Table S2. The resulting PCR products were cloned in pSDMA25 and validated by sequencing. The pSDMA25 vector containing the wild-type or catalytic mutant allele of Prp16 was linearized with restriction enzyme BaeI and introduced into the *PGAL7:Prp16* strain by biolistic transformation. NAT-resistant colonies were screened and validated by locus-specific PCR using primers listed in Supplementary Table S2. To generate strain that express U5 snRNA from safe haven locus the U5 locus along with 500bp upstream and 500bp downstream region, was first cloned in safe haven integration vector pSDMA57. The residues in loop 1 of U5 were mutated by inverse PCR using mutagenic primers in pSDMA57 vector.

### Media and growth conditions

Unless mentioned otherwise, for all experiments, strains were initially grown in permissive media YPG (2% peptone, 2% galactose, 1% yeast extract) overnight and then shifted to nonpermissive media YPD (2% peptone, 2% glucose, 1% yeast extract) for 12 hours. From these 12 hour YPD grown cultures, an aliquot of 0.1O.D. was subcultured into either permissive YPG media (for conditional overexpression of mCherry-Prp16) or nonpermissive YPD (for conditional knockdown of mCherry-Prp16) media and culture were grown for 9 hours. All strains were grown at 30°C.

### Confocal imaging and processing

To test the expression of mCherry-Prp16 under the *GAL7* promoter, both wild-type H99 and *PGAL7:*Prp16 strains were grown in permissive YPG media for overnight and then shifted to either permissive YPG or nonpermissive YPD media. The sample was collected every 3 hours for 12 hours of growth to observe the level of mCherry-Prp16 signal after the transfer into nonpermissive media. The cells were pelleted down, washed with PBS, and placed on an agarose patch contained in a microscopic slide. The coverslip was placed on the patch and was proceeded for imaging. The images were acquired in confocal laser scanning microscopy (Zeiss LSM 880) using a 63X oil immersion objective lens, and the images were processed by Zen Blue software and then analyzed by ImageJ.

### Semi quantitative RT-PCR and qRT-PCR

Total RNA was isolated from three independent batches of wild-type H99 and *PGAL7*:Prp16 cells grown in Prp16 knockdown condition (YPD), using triazole reagents and manufacturer protocol. The total RNA (20 μg) was subjected to DNaseI (NEB) digestion for 40 minutes as per manufacturer protocol. Reverse transcription was done on DNase-treated RNA using 10µM oligo dT, 10µM dNTPs and MMLV RT (NEB) according to the manufacture protocol. 100 ng of cDNA was amplified using primers specific to exon-intron-exon sequences. The PCR amplicons representing spliced and unspliced transcript segments were resolved on 8% native polyacrylamide gels. Signal intensities for the products were obtained by staining the gel with EtBr, followed by image acquisition using the Bio-Rad gel doc system. Quantification was done using ImageLab (V 6.1) and normalized to Actin control. For qRT PCR, the reactions were set up with 20-30ng of cDNA, 250 nM gene-specific primers, and FastStart Universal Sybr Green Master mix (Merck) in CFX Opus real-time system (BioRad). Fold change in transcript levels was calculated from the difference in cycle threshold value between Prp16 knockdown and wild type. To obtain the normalized threshold value (ΔΔCt), the ΔCt value was calculated by subtracting the Ct value for internal control – Actin, from the Ct value for each gene of interest. Then ΔΔCt was calculated by subtracting the wild type ΔCt value from the ΔCt value obtained for Prp16 knockdown. The fold change was calculated as 2^-(ΔΔCt). Primers used, and their sequences are given in Supplementary Table S2.

### RNA seq analysis

The DNase-treated and purified RNA samples were subjected to the next generation deep transcriptome sequencing. The sequencing service was outsourced to Molsys Pvt. Ltd Ahmadabad, following standard kits and protocols recommended by Illumina. Briefly, total mRNA was purified using Ribozero gold and the library was prepared with TruSeq Stranded Total RNA as per manufacturer protocols. The libraries were sequenced on the Illumina NovaSeq platform, which provided 150bp paired-end reads with 60-80 million reads. The raw files were subjected to quality check using fastQC. The adapters and low-quality reads were trimmed using fastP. The trimmed files were aligned to *C. neoformans* genome FungiDB-57 using STAR 2.7.3 with default parameters for twopassMode.

To analyze the genome-wide splicing alteration, the aligned (.bam) files from WT and Prp16 knockdown dataset were processed along with the gene annotation file (GTF) and genome sequence file (.fasta) by a Python script adapted from [20]. This scripted count reads mapped exon-intron junctions (EIJR), intron-exon junctions (IEJR) and exon-exon junctions (CSR). Introns with total read count < 10 were filtered out from the analysis. These counts were further imported to another python script [20] to calculate the intron retention level, the intron retention score (IRS), and the difference in IRS in Prp16 knockdown and WT (IRSdiff). As IRS values in WT and Prp16 knockdown follow a normal distribution, a two-sample Z-test was performed with the null hypothesis of zero difference in IRS between WT and Prp16 knockdown for each intron. These outputs were used to categorize the introns affected and unaffected classes for further analysis. To visualize intron retention for specific introns in WT and *PGAL7*:Prp16, sashimi plot was generated using python package rmats2Sashimi using aligned (.bam) files and gene annotation file (GTF) and genome sequence file (.fasta).

### Western blot

Wild-type and *PGAL7*:Prp16 cells were grown in permissive and nonpermissive media for 3-15 hours. The cells were pelleted and washed with PBS. Cell lysate was prepared using TCA method for western blot. Briefly cells were vortexed with acid-washed glass beads (Sigma, Cat. No. G8772) in 13% TCA for 30 mins at room temperature. The lysate was separated from the beads and pelleted at 13,000 rpm for 10 minutes and washed with 80% acetone to remove the residual TCA. The pellet was air-dried and resuspended in 4X Lamelli buffer (0.02% Bromophenol blue, 30% glycerol, 10% SDS, 250 mM Tris-Cl pH 6.8, 5% β-mercaptoethanol) and denatured at 95°C for 5 mins. The samples were loaded on 10% SDS PAGE, followed by electrophoresis, and transferred into PVDF membrane for 2 hours at 30 V using BioRad wet-transfer apparatus. The membrane was blocked with 3% BSA / 5% skimmed milk in TBS, depending on the primary antibody. After blocking, the membrane was incubated with primary antibody rabbit anti-mCherry (dilution 1:10000) (Abcam, Cat. No. ab213511) overnight at 4°C. The membrane was washed thrice with TBST (1X TBS + 0.1% tween20) and incubated with secondary antibody goat anti-rabbit HRP conjugated antibodies (dilution 1:10,000) for 1 hour at room temperature. The blot was washed thrice with TBST, and then the signals were then detected using the chemiluminescence method (Millipore Immobilon Forte HRP substrate).

### Visual data representation

The multiple sequence alignment was done using clustal omega. The scatter plots, CDF plots, bar plots and violin box plots from RNA seq data were generated using R studio (version 4.2.1) using the ggplot2 package. The growth curve and bar plot for quantification of semi-qRT-PCR was generated using GraphPad Prism (Version 8.4.2). Microsoft PowerPoint 2019 was used to design all the schematics and assemble the figures.

### Statistical and reproducibility

Unless mentioned otherwise, all the experiments were done in at least 3 batches. All the plots mention the standard deviation and the mean of independent experiments. The statistical significance was calculated with one-way ANOVA with Tukey’s multiple comparison tests or unpaired t-test. P valuesC≤ 0.05 were considered significant. All analyses were done using GraphPad prism version 8.4.2. The statistical significance of the plots generated from RNA seq data was calculated within the R package ggplot (equivalent to the Wilcoxon rank-sum test). The significances are mentioned as ns, P ≥ 0.05; * - P < 0.05; ** - P < 0.01; *** - P < 0.001; **** - P < 0.001.

## Author contributions

UV and MSN conceptualized and designed experiments for this study. MSN, VK and NS performed all experiments. MSN performed cloning, yeast strain generation, transcriptome and RT-PCR experiments. VK performed validatory qRT-PCR. NS performed transcriptomic data analysis and assisted MSN in site-directed mutagenesis. MSN performed detailed bioinformatics analysis and refined transcriptomic data with assistance from NS. MSN and UVR, interpreted the data and wrote the manuscript with contributions from VP.

## Supporting information

Supplementary Table S1. Strains used in this study

Supplementary Table S2. Sequences of the different primers and oligos used in this study.

Supplementary Table S3. Result of genome-wide splicing analysis from transcriptomic data.

## Acknowledgements

We acknowledge Prof. Kaustuv Sanyal, JNCASR, for the generous gift of the H99 wild-type strain and GAL7:mCherry vector. We also thank his lab members, including Dr. Vikas Yadav, PVS Satyadev, and Dr Shreyas Sridhar JNCASR, for their key suggestions during the course of this study. MolSys Labs Pvt. Ltd. Ahmadabad India. is acknowledged for RNA library preparation and deep sequencing service on the NovaSeq Illumina platform. The imaging facility at Department of MCB, IISc supported by DST-FIST program and Divisional Bioimaging facility, IISc are acknowledged.

## Supplementary information

**Supplementary Figure S1.** Multiple sequence alignment of C-terminal domain of predicted *C. neoformans* Prp16 (CNAG_02303) with Prp16 known in *S. cerevisiae, S. pombe, A. thaliana and H. sapiens*. Conserved motifs and residues used to generate mutants are highlighted.

**Supplementary Figure S2.** Validation of *C. neoformans* Prp16 conditional knockdown. (a) Schematics for growth conditions for validation of conditional knockdown of Prp16. (b) Semi-quantitative RT-PCR on RNA isolated from WT and *PGAL7*:Prp16 grown in YPD media for 12-15 hours. (c) Confocal imaging was used to examine the expression of mCherry-Prp16 under the *GAL7* promoter when WT and *PGAL7*:Prp16 cells were grown in YPG and YPD media for 12 hours. (d) Western blot for mCherry-Prp16 when *PGAL7*:Prp16 cells first grown in YPG media and shifted to YPD media for different time duration.

**Supplementary Figure S3.** (a) Schematic depiction of growth conditions for functionally significant knockdown of Prp16. (b) Conserved residues from helicase motif I and motif II chosen for generating Prp16 helicase domain mutants. (c) Growth profile by 10-fold serial dilution in agar plates incubated at 30 °C, 37°C and 16°C for WT, *PGAL7*:Prp16 and *PGAL7*:Prp16 strains expressing either wild type PRP16 (*PGAL7*:Prp16 SH:PRP16) or helicase domain mutants *PGAL7*:Prp16 SH:Prp16^K628A^ and *PGAL7*:Prp16 SH:Prp16 ^D719A^ from safe haven locus. (d) 5’ splice site and 3’ splice site residues of introns chosen for semi-quantitative RT-PCR. (e) Splicing assay of Cas35 intron7by semi-quantitative RT-PCR on RNA isolated from *C. neoformans* WT, *PGAL7*:Prp16 and *PGAL7*:Prp16 strains expressing WT CnPRP16 or the mutants K628A and D719A from safe haven locus. All strains were grown in the condition described previously for Prp16 knockdown. Quantification was done from three biological replicates. One-way ANOVA with Tukey’s multiple comparison test was used to calculate significance (ns, P ≥ 0.05; * - P < 0.05; ** - P < 0.01; *** - P < 0.001; **** - P < 0.001).

**Supplementary Figure S4.** (a) Normalized reads obtained from RNA seq of WT and *PGAL7*:Prp16, aligned to Prp16 locus. Decreased reads in *PGAL7*:Prp16 reaffirms transcriptional downregulation of Prp16 via promoter shutdown in YPD media. (b) and (c) Q-Q plot for IRS values in WT and *PGAL7*:Prp16, respectively. (d) In genome-wide splicing analysis, numbers of introns were found to be strongly affected, mildly affected, and unaffected. More than 88% of the introns are affected on Prp16 knockdown in *PGAL7*:Prp16 strain.

**Supplementary Figure S5.** (a) Violin-box plot of IRS in WT for each group of classified introns based on their last three nucleotides of 5’exon and 1^st^ nucleotide of 3’exon. (b) Violin-box plot of IRS differences between *PGAL7*:Prp16 and WT for each group of classified introns based on their last three nucleotides of 5’exon and 1^st^ nucleotide of 3’exon. Asterisks indicate statistically significant differences, as determined by the Wilcoxon rank-sum test within the R package ggplot2. (c) Schematic depiction of strain with GFP-Pac1E5I5E5 minigene under *H3* promoter at safe haven 3 locus. (d) Semi-quantitative RT-PCR for Pac1 E5I5E6 in WT, *PGAL7*:Prp16 strains with *PH3*:GFP-Pac1E5I5E6 integrated at safe haven 3 loci. RT-PCR was done with endogenous primers (which detect cDNA from both endogenous and minigene expressed transcripts) and minigene-specific primers. No amplicon from untransformed WT and *PGAL7*:Prp16 in the case of minigene-specific primers confirms accurate detection of mini-transcript expressed from safe haven locus.

**Supplementary Figure S6.** (a) Schematic depiction for pre-mRNA interaction with WT U5 and loop1 mutant U5^CUU-GAA^ for Pac1E5I5E6 and Mps1E6I6E7. (b) Schematic depiction of strain with WT U5 and loop1 mutant U5^CUU-GAA^ expressed from safe haven 1 locus. (c) CDF and (d) Violin-box plot of IRS differences between *PGAL7*:Prp16 and WT for each group of classified introns based on the size range of 10nts difference

**Supplementary Table S1.** Strains used in this study.

**Supplementary Table S2.** Sequences of the different primers and oligos used in this study.

**Supplementary Table S3.** Result of genome-wide splicing analysis from transcriptomic data.

