## Supplementary figures and images for "Prp16 enables efficient splicing of introns with diverse exonic consensus elements in the short-intron rich *Cryptococcus neoformans* transcriptome"

### Supplementary Figure S1

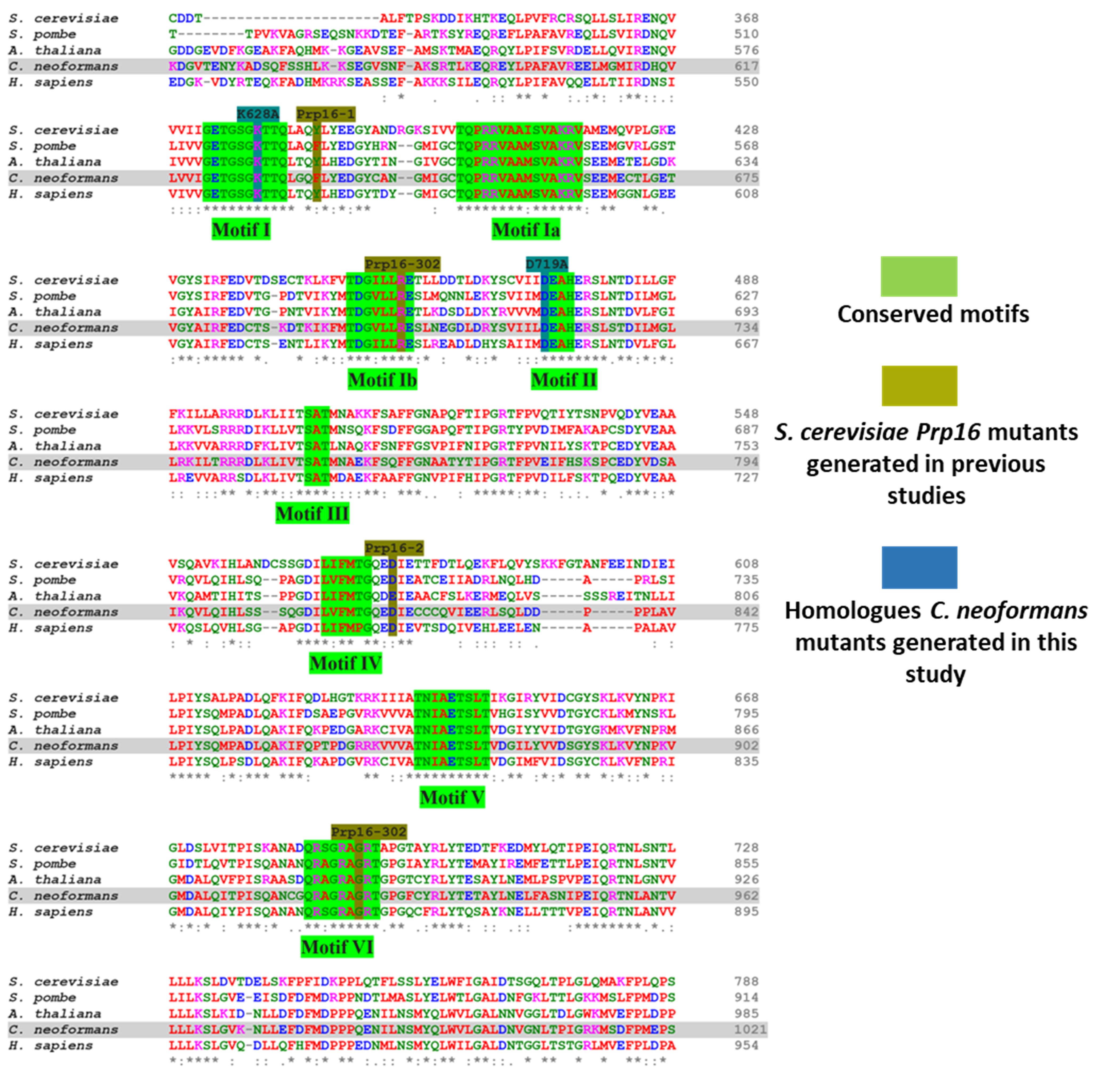

### Supplementary Figure S2

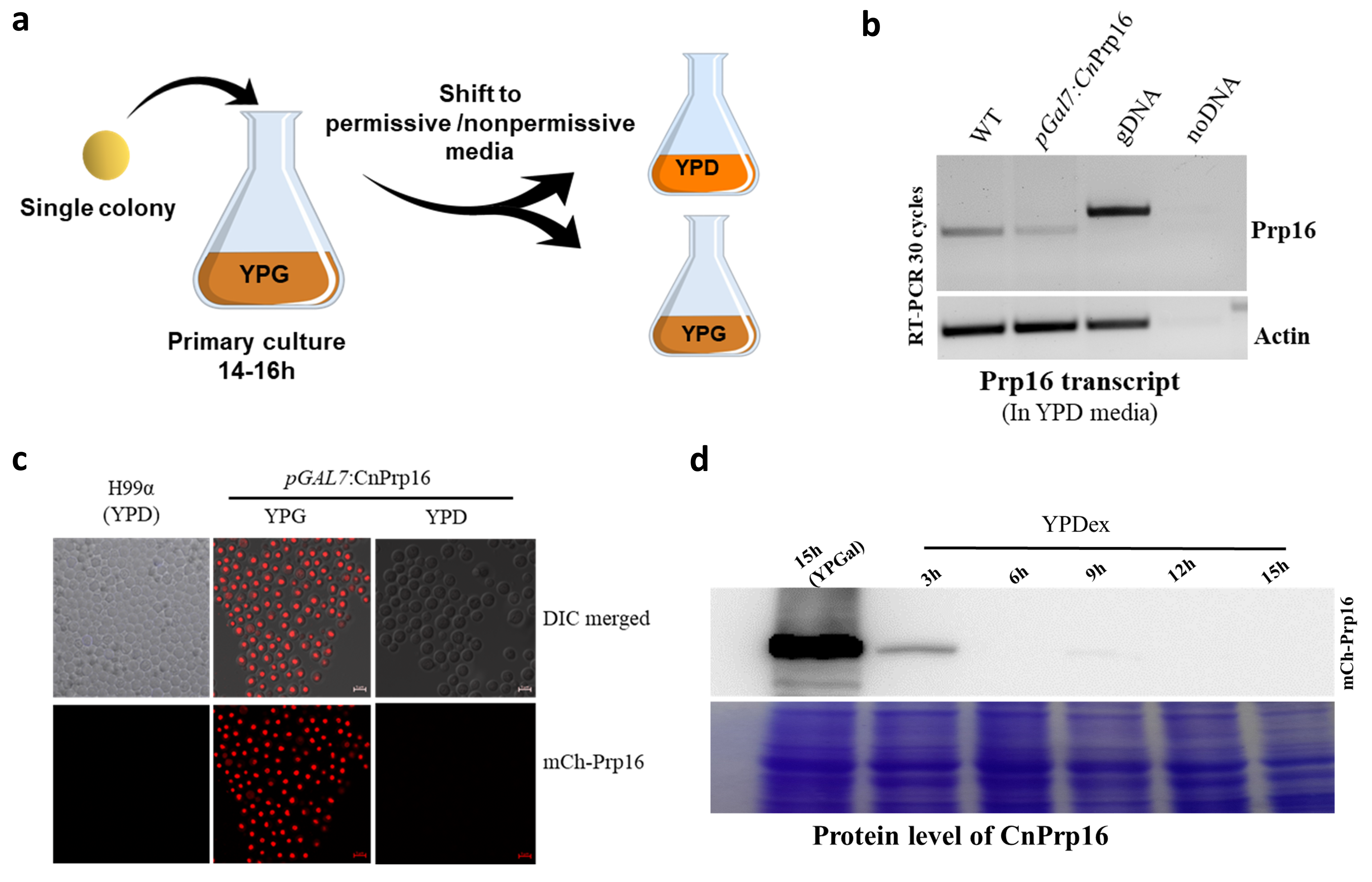

### Supplementary Figure S3

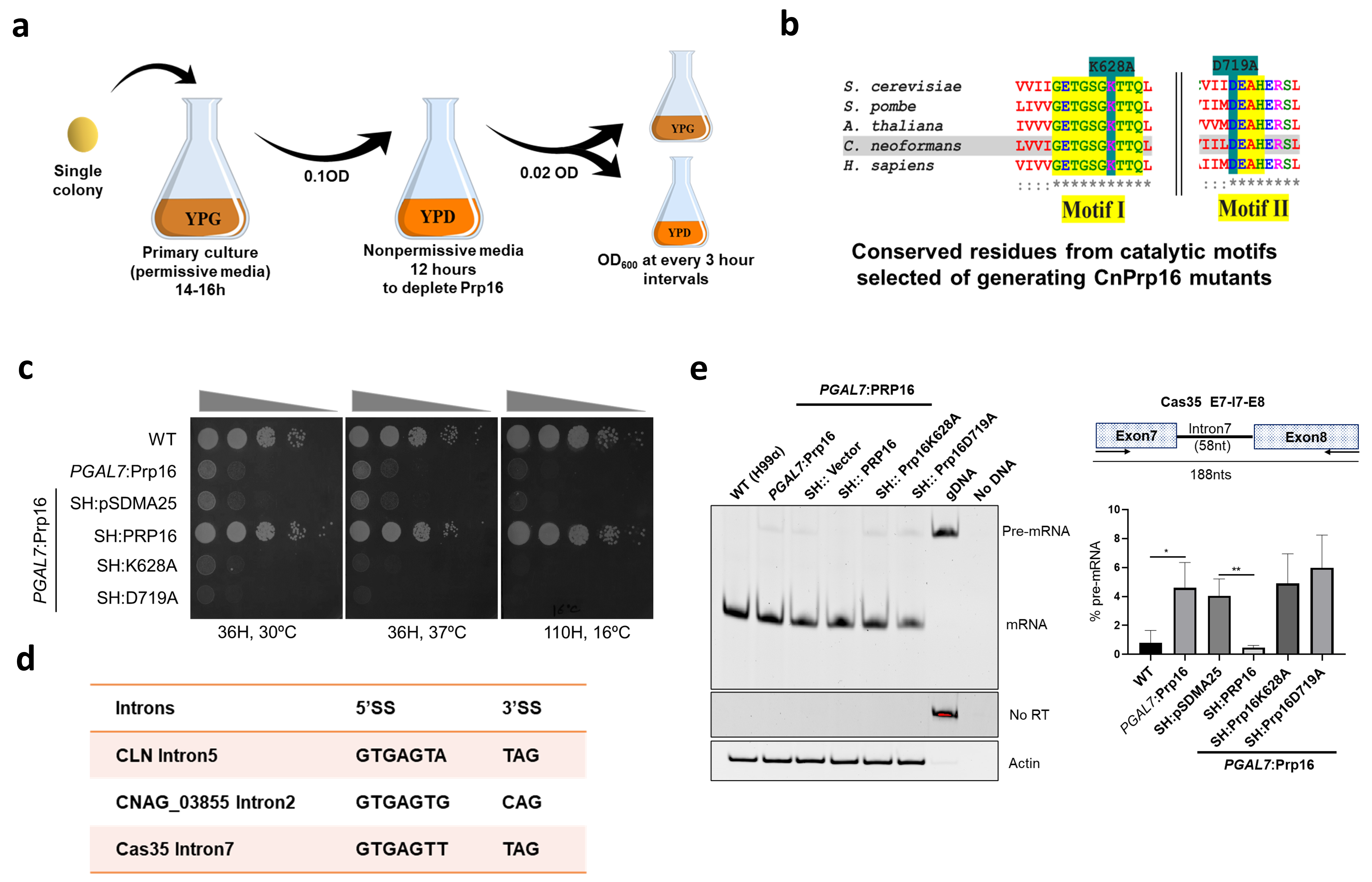

### Supplementary Figure S4

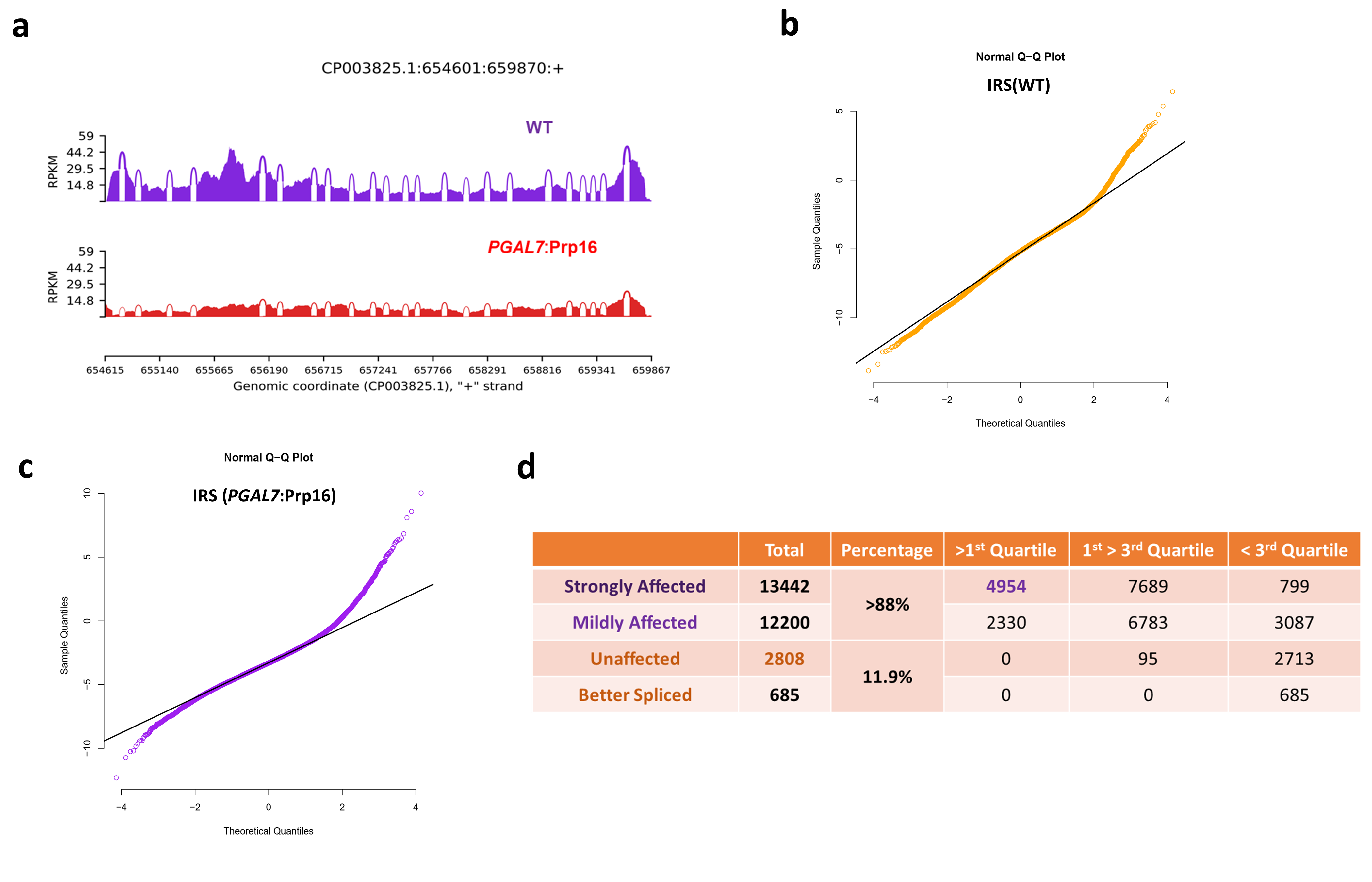

### Supplementary Figure S5

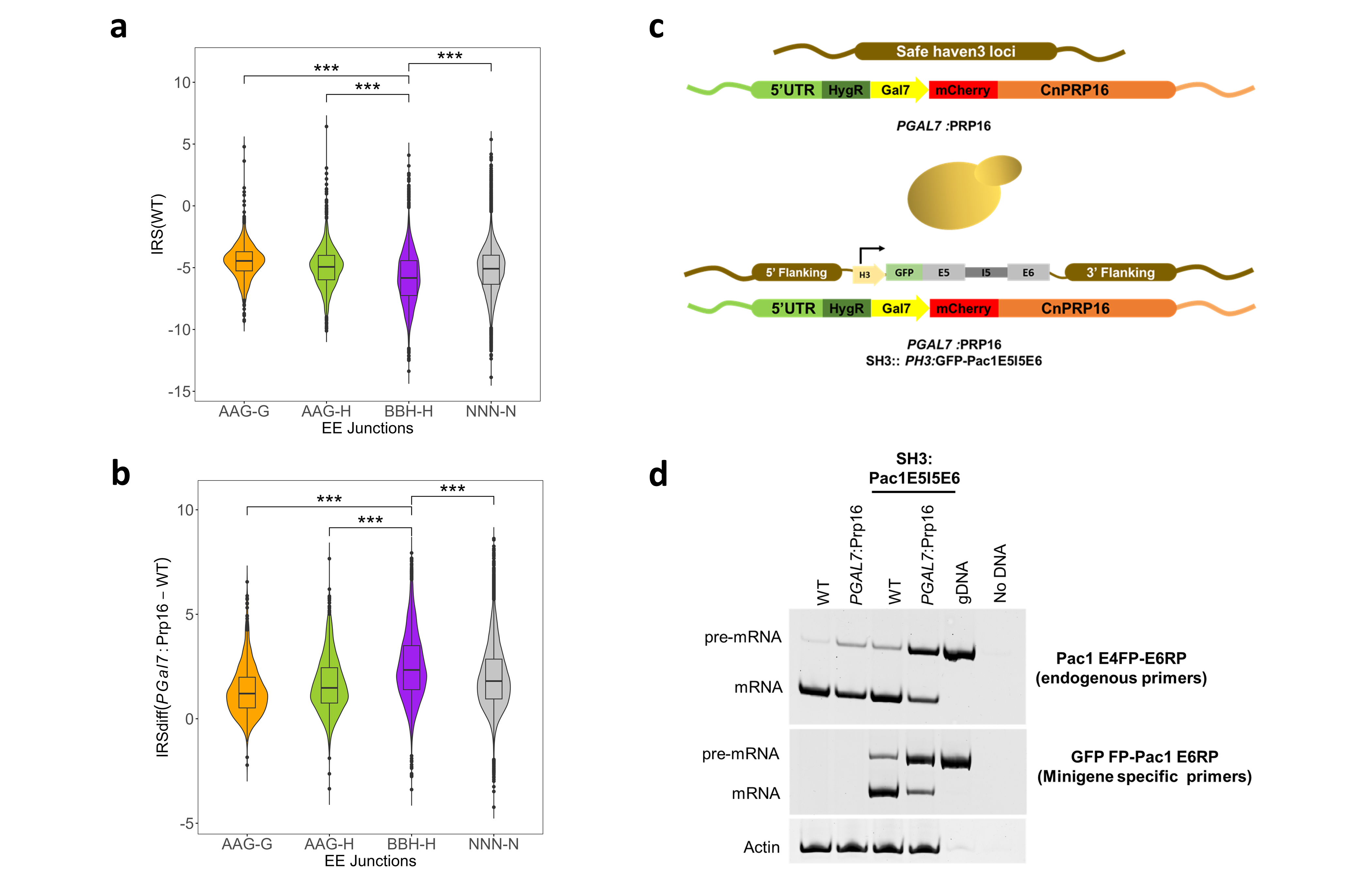

### Supplementary Figure S6

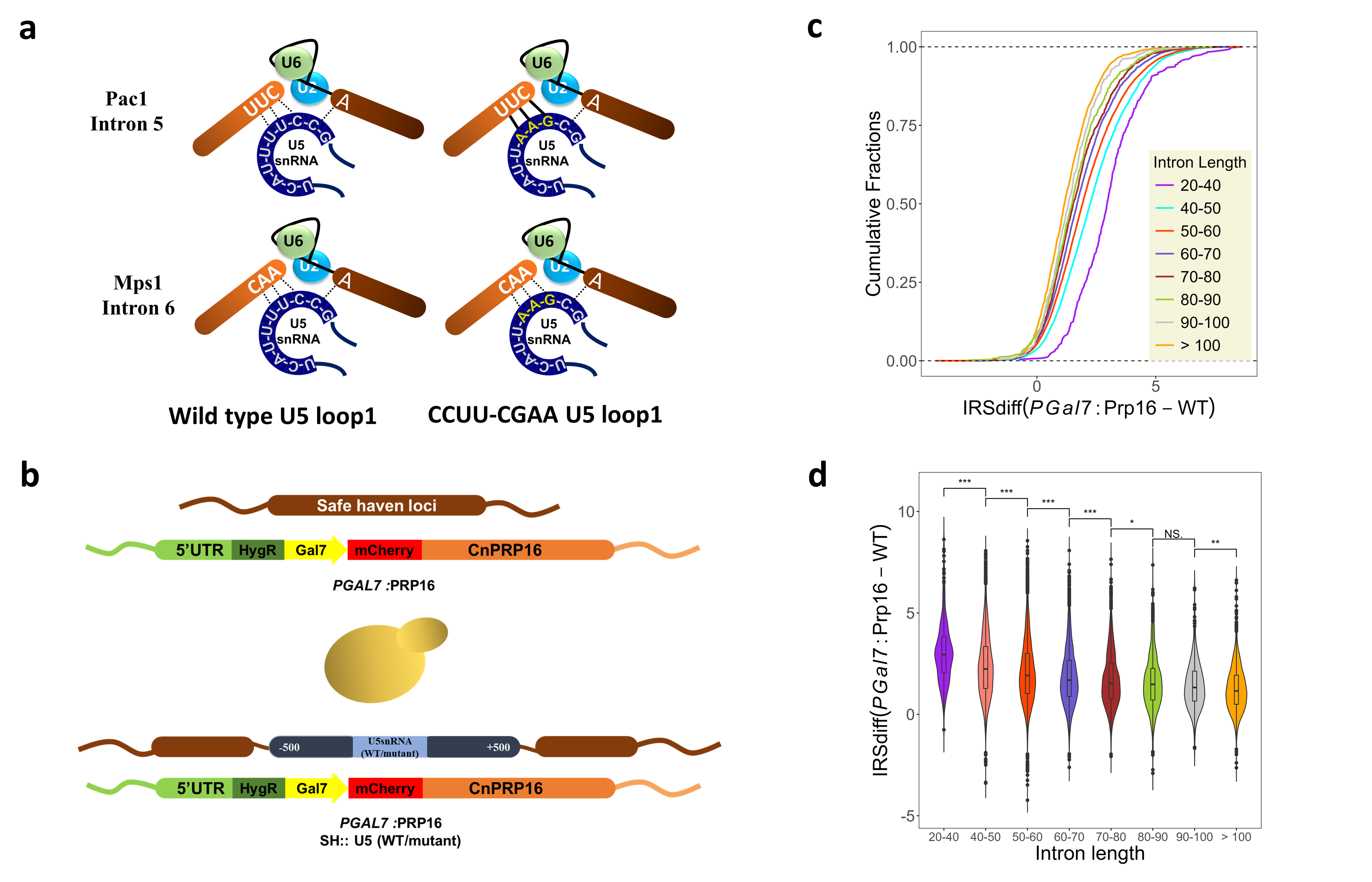
