## Supplementary Table S1. Strains used in this study for "Prp16 enables efficient splicing of introns with diverse exonic consensus elements in the short-intron rich *Cryptococcus neoformans* transcriptome"

*C. neoformans* Strains used in this study

| Strains | Genotype | Reference |
| --- | --- | --- |
| H99α | Wild type | ((Perfect et al., 1993)) |
| *PGAL7*:Prp16 | MATα, PRP16:: HygB-GAL7p-mCherry-Prp16 | This study |
| *PGAL7*:Prp16, SH:PRP16 | MATα, PRP16:: HygB-*PGAL7*-mCherry:Prp16, SH::PRP16-NAT (pSDMA25) | This study |
| *PGAL7*:Prp16, SH::K628A | MATα, PRP16:: HygB-*PGAL7*-mCherry:Prp16, SH::Prp16 K628A-NAT (pSDMA25) | This study |
| *PGAL7*:Prp16,  SH:: D719A | MATα, PRP16::HygB-GAL7p-mCherry-Prp16, SH::PRP16 D719A-NAT (pSDMA25) | This study |
| H99α, SH3::Pac1I5 | MATα, SH3:: *PH3*-GFP:Pac1E5-I5-E6-NAT (pEE27) | This study |
| H99α, SH3::Pac1I5 AAG | MATα, SH3:: *PH3*-GFP:Pac1E5-I5-E6(5’SS TTC-AAG)-NAT (pEE27) | This study |
| H99α, SH3::Pac1I5  AAG-G | MATα, SH3:: *PH3*-GFP:Pac1E5-I5-E6(5’SS-3’SS TTCA-AAGG)-NAT (pEE27) | This study |
| *PGAL7*:Prp16, SH::Pac1I5 | MATα, Prp16:: HygB-GAL7p-mCherry-Prp16, SH3:: *PH3*-GFP:Pac1E5-I5-E6-NAT (pEE27) | This study |
| *PGAL7*:Prp16,  SH::Pac1I5 AAG | MATα, Prp16:: HygB-GAL7p-mCherry-Prp16, SH3:: *PH3*-GFP:Pac1E5-I5-E6(5’SS TTC-AAG)-NAT (pEE27) | This study |
| *PGAL7*:Prp16,  SH::Pac1I5 AAG-G | MATα, Prp16:: HygB-GAL7p-mCherry-Prp16, SH3:: *PH3*-GFP:Pac1E5-I5-E6(5’SS,3’SS TTCA-AAGG)-NAT (pEE27) | This study |
| *PGAL7*:Prp16,  SH::U5 | MATα, PRP16:: HygB-*PGAL7*-mCherry:Prp16, SH::U5 -NAT (pSDMA57) | This study |
| *PGAL7*:Prp16,  SH::U5 CUU-GAA | MATα, PRP16:: HygB-*PGAL7*-mCherry:Prp16, SH:: U5 CUU-GAA -NAT (pSDMA57) | This study |

Perfect, J. R., Ketabchi, N., Cox, G. M., Ingram, C. W., & Beiser, C. L. (1993). Karyotyping of Cryptococcus neoformans as an epidemiological tool. *Journal of Clinical Microbiology*, *31*(12), 3305–3309. <https://doi.org/10.1128/jcm.31.12.3305-3309.1993>
