## Supplementary Table S2. Sequences of the different primers and oligos used in this study. for "Prp16 enables efficient splicing of introns with diverse exonic consensus elements in the short-intron rich *Cryptococcus neoformans* transcriptome"

Primers/oligos used in this study

| **SN** | **Primer name** | **Sequence** | **Description** |
| --- | --- | --- | --- |
| **1** | Prp16 H99 5’UTR FP SACI | TCA GAG CTC TGATTCTCTTCATTCTCCT | Amplification of 1kb upstream fragment from ATG of *C. neoformans* Prp16 |
| **2** | Prp16 H99 5’UTR RP SACI | TCAGAGCTCGTTCAGGTATGTAGAGGATGA |  |
| **3** | H99 PRP16 P1 HINDIII | TCCAAGCTTATGTCCGCCAGGTCACCAA | Amplification of 1kb downstream fragment from ATG of *C. neoformans* Prp16 |
| **4** | H99 PRP16 P2 XHOI | ATGCTCGAGTTCATTCCATCTGTCCCTGTTT |  |
| **5** | mCherry FP | AACATCAAGTTGGACATCACCTCCCA | mCherry FP for locus specific PCR validation of Gal7:mCherry-Prp16 |
| **6** | Prp16 (4350) RP | ACTCTTTAGCGTTGAGATCC | Used with mCherry FP for locus specific PCR validation of Gal7:mCherry-Prp16 |
| **7** | Prp16 (176 CDS) RP | GTCGAAACAGCTCTAAGGAATGC | Used with H99 PRP16 P1 HINDIII for qRT-PCR of Prp16 |
| **8** | CnH99Prp16RP (XbaI) | TTGTCTAGATTAAATGCCTCCTGCTCGTC | Used with H99 PRP16 P1 HINDIII for amplification of full length gene body of Prp16 or cDNA |
| **9** | CnPrp16-500FP(XhoI) | GTACCGGGCCCCCCCTCGAGAACGTAGAATACCGCGAAATCG | Used to clone Prp16 loci in safe haven plasmid |
| **10** | CnPrp16+500RP(PstI) | GTGGATCCCCCGGGCTGCAGATTGAAGAAGAACCAGTTCAGC |  |
| **11** | cnPrp16_K628A_FP | GGGTCAGGCGCAACAACTCAG | For K628A site directed mutagenesis of Prp16 |
| **12** | cnPrp16_K628A_RP | CTGAGTTGTTGCGCCTGACCC |  |
| **13** | cnPrp16_D719A_FP | ATCATCCTTGCTGAAGCACAC | For D719A site directed mutagenesis of Prp16 |
| **14** | cnPrp16_D719A_RP | GTGTGCTTCAGCAAGGATGAT |  |
| **15** | MPS1I6_FP | CGTTCATACAGATTTGAAGCC | RT-PCR OF Mps1 E6I6E7 |
| **16** | MPS1I6_RP | CTTCAATACCTTCTGATTGTTC |  |
| **17** | 3855I2_FP | GGAACTCTCTACGAGATGGG | RT-PCR OF CNAG_03855 E2I2E3 |
| **18** | 3855I2_RP | GTATGGCGAAAAGTTGTGTCG |  |
| **20** | SEC72I4_FP | CAATACGAGGACGCAAAGC | RT-PCR OF Sec72 E4I4E5 |
| **21** | SEC72I4_RP | TCATTGAACGGTGTCTGGG |  |
| **22** | CAS35I7_FP | AGAGGTGGAGGATGTTATTGC | RT-PCR OF Cas35 E7I7E8 |
| **23** | CAS35I7_RP | ATTCCAAGAATCTTGAAAGGCG |  |
| **24** | 2654I4_FP | GACGAAAATAGAGCATGGGG | RT-PCR OF DNA ploξ E4I4E5 |
| **25** | 2654I4_RP | TCATAAGCCATAACGACTGG |  |
| **26** | 00649I3_FP | GCCGACATCATCGAACTCG | RT-PCR OF CNAG_00649 E3I3E4 |
| **27** | 00649I3_RP | CCATGAACAAGACGGGAGC |  |
| **29** | RPC1I11_FP | TGGAGGTATCGTGCAGTTCC | RT-PCR OF RPC1 E11-I11-E12 |
| **30** | RPC1I11_RP | TGGGAGGAACAAAAGACTCG |  |
| **31** | SH FP1 | GGGTATGCCACAGATGCAGAT | Used for validation of integration at safe haven loci |
| **32** | SH RP1 | ACTGGTGAGTACTCAACCAAG |  |
| **33** | SH FP2 | TCAGCAACGCCGTTGAATCCT |  |
| **34** | SH RP2 | TTGGATCCTCAATTGTCTCCT |  |
| **35** | SH3FP | TCTACGTTGGCGCTTCAAGC | Used with SH FP2 and SH RP1 for validation of integration at safe haven 3 loci |
| **36** | SH3RP | TTGGAGTCAACAGCCGTGGG |  |
| **37** | U5_up_FP | CTGCAGTCTGCTGACGAACTCTATTC | Used for amplification and cloning of U5 snRNA with 500bp upstream and downstream sequence |
| **38** | U5_down_RP | CTCGAGGTACAGGAATAGACTCGTCG |  |
| **39** | Pac1I5(5PSS-AAG)FP | CTACGGACAAAGGTACGTCGT | 5’SS TTC-AAG mutagenesis of pac1 E5I5E6  3’SS C-G mutagenesis of pac1 E5I5E6 |
| **40** | Pac1I5(5PSS-AAG)RP | ACGACGTACCTTTGTCCGTAG |  |
| **41** | Pac1I5(5PSS-AAG)FP | CTACGGACAAAGGTACGTCGT |  |
| **42** | Pac1I5(5PSS-AAG)RP | ACGACGTACCTTTGTCCGTAG |  |
| **43** | U5-CUU_FP | CGAATAAATCTCTCGCGAATTACTAGAGATATCC | Used for Site directed mutagenesis of U5 loop1 CCU-GGA |
| **44** | U5-CUU_RP | GGATATCTCTAGTAATTCGCGAGAGATTTATTCG |  |
